# Genotypic and phenotypic diversity within the neonatal HSV-2 population

**DOI:** 10.1101/262055

**Authors:** Lisa N. Akhtar, Christopher D. Bowen, Daniel W. Renner, Utsav Pandey, Ashley N. Della Fera, David W. Kimberlin, Mark N. Prichard, Richard J. Whitley, Matthew D. Weitzman, Moriah L. Szpara

**Affiliations:** Department of Pediatrics, Division of Infectious Diseases, Children’s Hospital of Philadelphia; Department of Biochemistry and Molecular Biology, Center for Infectious Disease Dynamics, and the Huck Institutes of the Life Sciences, Pennsylvania State University; Department of Pathology and Laboratory Medicine, Division of Protective Immunity and Division of Cancer Pathobiology, Children’s Hospital of Philadelphia; Department of Pediatrics, Division of Infectious Diseases, University of Alabama at Birmingham; University of Pennsylvania Perelman School of Medicine

**Keywords:** Neonatal, herpes simplex virus 2, human herpesvirus 2, comparative genomics, viral spread, minor variants

## Abstract

More than 14,000 neonates are infected with herpes simplex virus (HSV) annually. Approximately half display manifestations limited to the skin, eyes, or mouth (SEM disease). The rest develop invasive infections that spread to the central nervous system (CNS disease or encephalitis) or systemically (disseminated disease). Invasive HSV disease is associated with significant morbidity and mortality, but viral and host factors that predispose neonates to these forms are unknown. To define viral diversity within the infected neonatal population, we evaluated ten HSV-2 isolates from newborns with a range of clinical presentations. To assess viral fitness independent of host immune factors, we measured viral growth characteristics in cultured cells and found diverse *in vitro* phenotypes. Isolates from neonates with CNS disease were associated with larger plaque size and enhanced spread, with isolates from cerebrospinal fluid (CSF) exhibiting the most robust growth. We sequenced complete viral genomes of all ten neonatal viruses, providing new insights into HSV-2 genomic diversity in this clinical setting. We found extensive inter-host and intra-host genomic diversity throughout the viral genome, including amino acid differences in more than 90% of the viral proteome. The genes encoding glycoprotein G (gG, US4), gI (US7), gK (UL53), and viral proteins UL8, UL20, UL24, and US2 contained variants that were found in association with CNS isolates. Many of these viral proteins are known to contribute to cell spread and neurovirulence in mouse models of CNS disease. This study represents the first application of comparative pathogen genomics to neonatal HSV disease.

**Importance:** Herpes simplex virus (HSV) causes invasive disease in half of infected neonates, resulting in significant mortality and permanent cognitive morbidity. The factors that contribute to invasive disease are not understood. This study reveals diversity among HSV isolates from infected neonates, and makes the first associations between viral genetic variations and clinical disease manifestations. We found that viruses isolated from newborns with encephalitis show enhanced spread in culture. These viruses contain protein-coding variations not found in viruses causing non-invasive disease. Many of these variations are found in proteins known to impact neurovirulence and viral spread between cells. This work advances our understanding of HSV diversity in the neonatal population and how it may impact disease outcome.

## Introduction

Each year an estimated 10,000 neonates are infected with HSV-2, and 4,000 infected with HSV-1, worldwide (1). Infants are typically infected at the time of birth by maternal genital shedding of HSV, most often by mothers who are not aware of their infection (2–4). The recent increase in genital HSV-1 incidence among women of childbearing age, particularly in developed nations, suggests that the burden of neonatal infection will continue to rise (1, 5). While some infected infants exhibit only superficial infection limited to the skin, eyes, or mouth (SEM disease; 45%), about half develop invasive systemic (disseminated disease; 25%) or central nervous system (CNS disease; 30%) infections associated with significant morbidity and mortality (6, 7). Currently, the antiviral medication acyclovir is the standard therapy for all forms of neonatal HSV disease. Although this intervention has reduced mortality due to invasive disease, most survivors of invasive disease are left with permanent neurodevelopmental deficits (8, 9).

The factors that predispose a neonate to invasive HSV infection are not entirely known. Recent studies have found that some adults and children outside of the neonatal period who experience HSV infection of the brain have a host genetic defect within the Toll-like receptor-3 (TLR3) pathway (10, 11). Outside of the neonatal period, HSV encephalitis is rare, as are host defects in the TLR3-pathway. By contrast, half of HSV-infected neonates experience invasive CNS or disseminated disease, making it less likely that host genetic defects alone could account for all of the observed cases of invasive infection in neonates. Prior clinical data on mother-to-infant transmission of HSV indicate that most cases of neonatal disease, including invasive forms of disease, result from newly acquired or primary HSV infection prior to the development of maternal antibody production (2–4, 12). This suggests a window of opportunity where the contributions of viral genetic variation to the progression of invasive infection and disease may be greater than in adults.

Prior studies have identified viral genetic factors that influence virulence or disease for reoviruses, influenza virus, HIV, and others (13–18). In contrast to these RNA viruses, HSV was presumed to have lower genetic diversity and potential for variation in virulence, due to its relatively stable DNA genome and long co-evolutionary history with humans (19). The assumption of limited HSV heterogeneity was supported by early studies that utilized low-resolution restriction fragment length polymorphism (RFLP) or single-gene analyses to compare multiple HSV isolates (20–22). However Rosenthal and colleagues used RFLP and PCR analysis of a single locus to demonstrate that a heterogeneous HSV population can exist in an invasive neonatal infection, and provided proof of principle that natural genetic variation can impact neurovirulence (23, 24). More recently, advances in high-throughput sequencing (HTSeq) have enabled a re-evaluation of herpesvirus genome-wide variation, which suggests that herpesviruses harbor extensive diversity both between strains or individuals (inter-host variation) as well as within a single individual (intra-host variation) (25–27). These minor genetic variants may become clinically important if a variant within the viral population becomes the new dominant allele or genotype as a result of a bottleneck at transmission, entry into a new body compartment, or selective pressure such as antiviral therapy (28, 29).

Several recent examples have demonstrated the potential for new insights gained by applying HTSeq approaches to herpesvirus infections in a clinical setting. In HTSeq studies of congenital infection by the beta-herpesvirus human cytomegalovirus (HCMV), Renzette *et al.* found evidence for heterogeneous viral populations both within and between hosts (30–35). The levels of diversity observed in congenital HCMV infections far exceeded those observed in adult infections (36–38). HTSeq-based examination of vaccine-associated rashes due to the alpha-herpesvirus varicella zoster virus (VZV) demonstrated that adult skin vesicles contain a subset of the viral population introduced during vaccination, and found at least 11 VZV genomic loci that were linked to rash formation (26). Recent HTSeq comparisons of adult genital HSV-2 revealed the first evidence of a single individual shedding two distinct strains (39), demonstrated changes in the viral genome over time in a recently infected host (40), and provided the first evidence of ancient recombination between HSV-1 and HSV-2 (41, 42). However, to date there has been no evaluation of genome-wide variation in neonatal HSV isolates to determine the levels of diversity in this population, or the potential impact(s) of viral genetic variants on disease.

Until recently, several technical barriers prevented thorough assessment of neonatal HSV genomes. A key constraint on studies of neonatal disease has been the availability of cultured, minimally-passaged viral samples that are both associated with clinical information and have also been maintained in a low-passage state that is appropriate for sequencing and further experimental studies. Historically viral culture was part of the HSV diagnostic workflow, but this has been superseded in clinical laboratory settings by the speed and sensitivity of viral detection by PCR (6, 43, 44). This change limits neonatal HSV sample availability for *in vitro* and animal model studies. In addition, many previously archived neonatal HSV isolates have been passaged extensively, allowing them to acquire mutations that enhance viral growth in culture (23, 24). Additional challenges for HTSeq approaches to neonatal HSV include the large size of the viral genome (∼152 kb), its high G + C content (∼70%), and a large number of variable-number tandem repeats in the viral genome (> 240 mini/micro-satellite repeats and >660 homopolymers of ≥ 6 base pairs) (27, 45). Therefore, many studies of HSV diversity (21, 46–49) or the effect of HSV genetic variation on disease (50–52) have relied on low-resolution restriction fragment length polymorphism (RFLP) or single-gene PCR analyses, due to the speed and ease of analysis in comparison to whole-genome approaches (53–58). To overcome these challenges, we combined our expertise in HSV comparative genomics and phenotypic analysis (53, 58–60) with a unique resource of low-passage, well-annotated neonatal specimens (8, 9, 61).

Here we analyzed a set of ten low-passage clinical HSV-2 isolates collected from neonates with HSV infection, enrolled in one of two clinical studies that spanned three decades of patient enrollment (1981-2008) (8, 9, 61). These samples represented a wide range of clinical manifestations including SEM, CNS, and disseminated disease, and each sample was associated with de-identified clinical information. We defined the level of diversity in this population using comparative genomics and an array of cell-based phenotypic assays. We found that HSV-2 isolates displayed diverse *in vitro* phenotypes, as well as extensive inter- and intra-host diversity distributed throughout the HSV-2 genome. Finally, we found coding variations in several HSV-2 proteins associated with CNS disease. This study represents the first-ever application of comparative pathogen genomics to neonatal HSV disease and provides a basis for further exploration of genotype-phenotype links in this clinically vulnerable patient population.

## Results

### Neonatal HSV-2 samples represent a diverse clinical population

We utilized samples collected from ten HSV-2-infected neonates enrolled by the National Institute of Allergy and Infectious Diseases Collaborative Antiviral Study Group (CASG) for clinical trials between 1981 and 2008 (8, 9, 61). These infants encompassed a range of clinical disease manifestations (see Table 1), with about half experiencing invasive CNS disease (5 patients) or disseminated (DISS) disease with CNS involvement (2 patients), and the remainder experiencing non-invasive SEM disease (3 patients). Extensive clinical information was available for each patient, including long-term neurocognitive and motor outcomes (Table 1). This population was also diverse with respect to sex, race, gestational age, and enrollment center (Table 1; enrollment center data not shown). All samples were collected at the time of diagnosis, prior to initiation of acyclovir therapy. Each isolate was cultured once as part of the diagnostic process, with expansion only for the experiments shown here. Although the sample size was constrained by the rarity of neonatal HSV infection and availability of appropriately maintained isolates, our group is similar in size to prior HTSeq comparisons of congenital HCMV samples (30–35), and is the largest group of neonatal HSV samples ever subjected to comparative genomic and phenotypic analysis.

**Table 1.** Clinical characteristics associated with HSV-2 isolates from ten patients.

| Clinical Isolate* | Clinical Disease at Diagnosis | Sample Source | Morbidity Score - Mental | Morbidity Score - Motor | Patient Age at Disease Onset (days) | Gestational Age at Time of Birth (weeks) | Patient Sex & Race |
| --- | --- | --- | --- | --- | --- | --- | --- |
| **CNS11** | CNS | CSF | 4 | 4 | 12 | 37 | M, W |
| **DISS14** | DISS + CNS | CSF | 2 | 2 | 7 | 39 | M, W |
| **CNS03** | CNS | SKIN | 4 | 4 | 17 | 37 | M, W |
| **CNS15** | CNS | SKIN | 3 | 4 | 19 | 36 | M, W |
| **CNS17** | CNS | SKIN | 4 | 4 | 17 | 40 | M, W |
| **DISS29** | DISS + CNS | SKIN | 3 | 3 | 5 | 38 | F, B |
| **CNS12** | CNS | SKIN | 4 | 4 | 16 | 41 | F, W |
| **SEM02** | SEM | SKIN | 1 | 1 | 5 | 38 | F, W |
| **SEM13** | SEM | SKIN | 4 | 4 | 11 | 27 | F, B |
| **SEM18** | SEM | SKIN | 2 | 2 | 17 | 37 | F, W |

\* Clinical isolate order based on **Figure 1**.

### Neonatal HSV-2 isolates have different fitness in culture

To determine whether the viruses isolated from this neonatal population (Table 1) were intrinsically different, we assessed viral growth in culture, which provides a consistent environment that is independent of host genetic variation. To minimize the impact of immune pressure we selected Vero monkey kidney cells, which lack an interferon response (62, 63). Each viral isolate was applied to a confluent monolayer of cells *in vitro*, and allowed to form plaques for 100 hours (h) (Figure 1A). Average plaque size differed between isolates, with six of the ten isolates being statistically larger (indicated in green) than the other four (indicated in black, Figure 1B) (one-way ANOVA with Holm-Sidak’s multiple comparisons test, p < 0.05). The average plaque size of the previously described low passage, adult HSV-2 isolate SD90e is shown for comparison (64). The largest plaque sizes were observed in the two viruses isolated directly from the cerebrospinal fluid (CSF; isolates CNS11 and DISS14; Figure 1B); these two isolates were not statistically different from one another in size, but were significantly larger than any of those isolated from the skin (one-way ANOVA with Holm-Sidak’s multiple comparisons test, p < 0.05). Plaque size was assessed at each passage in culture and remained constant from the time the isolates were received in our laboratory (passage 2) through their genetic and phenotypic analysis (passage 4). The variance in plaque sizes produced by a given isolate was not statistically different between isolates (Figure 1B). The differences in average plaque size between isolates suggested that the HSV-2 populations found in each neonatal isolate are indeed intrinsically different.

**Figure 1.**
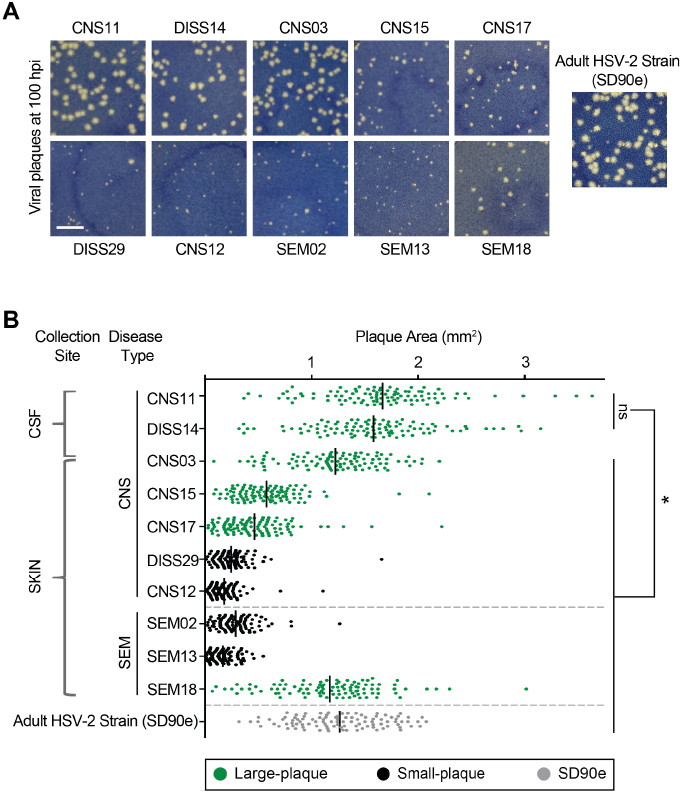
Neonatal HSV-2 isolates generate different sized plaques in culture. (**A**) Representative plaques are shown after virus incubation on Vero cells for 100 h. The previously described low-passage adult HSV-2 strain SD90e is shown for comparison (64). Scale bar = 5 mm. (**B**) Quantification of plaque area on Vero cells. Dots represent 100 individually measured plaques and black bar represents the mean. Each green isolate (large-plaque) is statistically larger than each black isolate (small-plaque). Black isolates are not statistically different from one another. Additionally, each CSF-derived isolate is statistically larger than all other isolates shown. For all statistics, p < 0.05 by one-way ANOVA followed by Holm-Sidak’s multiple comparisons test.

### Entry kinetics, DNA replication, protein expression, and virus production do not account for differences in plaque size

Plaque formation is a complex endpoint that involves the ability of the virus to enter the cell, replicate its double stranded DNA genome, produce viral proteins and assemble new virions that spread to adjacent cells. Therefore, we explored whether the differences in plaque formation observed in Vero cells reflected inherent differences in the ability of isolates to complete each of these stages of the viral life cycle. For these comparisons, two large-plaque-forming isolates (CNS11 and CNS03) were compared to two small-plaque-forming isolates (CNS12 and SEM02). First, we compared each isolate’s rate of cell entry. Virus was applied to chilled cells, followed by warming to synchronize cell entry (Figure 2A). A low pH solution was applied at various points over the first hour of cell entry to inactivate any virus that had not yet entered a cell, and plaque formation was then allowed to proceed for 100 h. We found no difference in rates of cell entry between these four representative viral isolates (Figure 2B), suggesting that large plaques did not result from increased rates of virus entry into cells.

**Figure 2.**
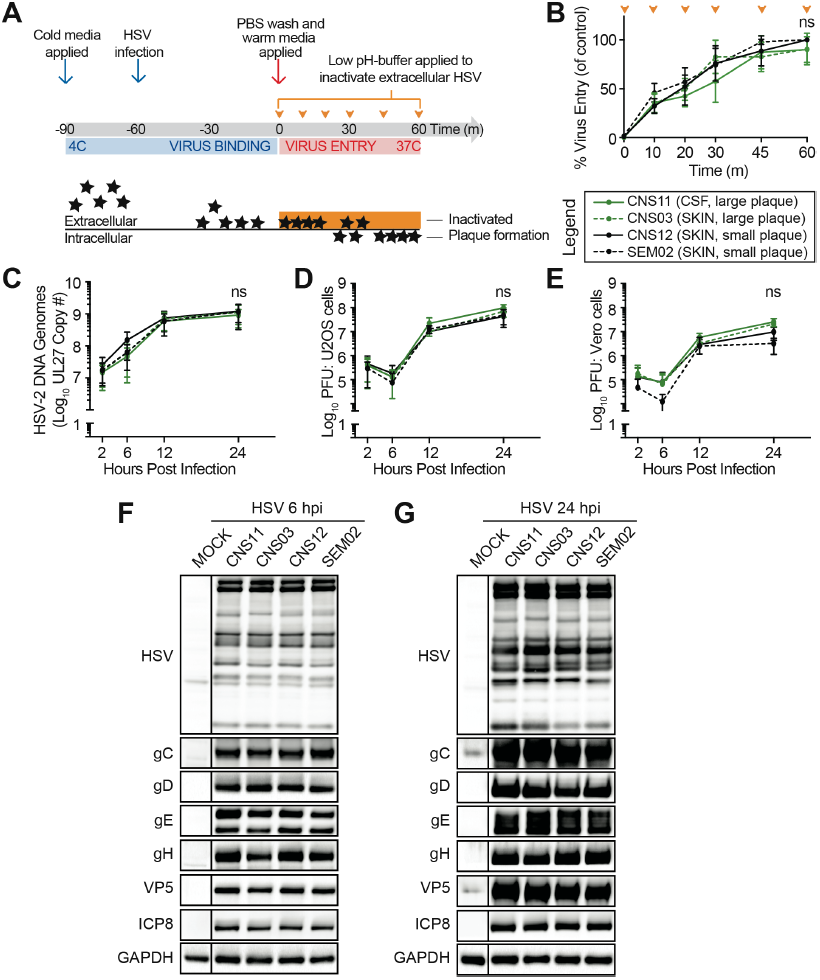
Increased plaque size in culture is not determined by viral entry, DNA replication, protein expression, or infectious virus production. Viral growth characteristics were compared for representative neonatal isolates, including large-plaque-formers (green) and small-plaque-formers (black). **(A-B)** Viral entry kinetics. **(A)** Viral isolates were applied to Vero cell monolayers at 4°C for 1 h to allow virus binding, then moved to 37°C to allow virus entry. Extracellular virus was inactivated by a low-pH buffer at the times indicated (orange arrowheads). Cell monolayers were washed and overlaid with methylcellulose. Plaques were scored after 100 h of incubation. **(B)** Viral entry was quantified as the fraction of plaques formed following citrate buffer application, where 100% is the number of plaques formed on a monolayer not treated with citrate buffer (control). These data represent three independent experiments. Two-way ANOVA followed by Tukey’s multiple comparison test was applied. **(C-E)** Single-cycle viral replication kinetics. Vero cell monolayers were infected at MOI = 5 and incubated in the presence of 0.1% human serum. Cell monolayers were harvested at the time points indicated. **(C)** The quantity of viral genomes present was evaluated by qPCR for UL27. Infectious virion production (titer) was evaluated by plaque formation on U2OS **(D)** or Vero **(E)** cells. These data represent three independent experiments. Two-way ANOVA followed by Tukey’s multiple comparison test was applied. **(F-G)** Protein production. Vero cell monolayers were infected at MOI = 5 for 6 h **(F)** or 24 h **(G)**. Whole cell lysates were subjected to immunoblot analysis with the following antibodies: gC/gD/gE/gH, four virion glycoproteins; VP5 (UL19), capsid protein; ICP8 (UL29), viral single-strand DNA-binding protein; HSV, viral antibody against whole HSV-1; and GAPDH, cellular glyceraldehyde-3 phosphate dehydrogenase as a loading control.

We next infected Vero cells at high multiplicity of infection (MOI = 5) to compare the outcome of a single round of viral replication. We found that all four isolates produced similar numbers of genome copies (as measured by qPCR for the gB gene; see Methods for details) (Figure 2C), indicating that differences in viral DNA replication did not influence plaque size. We quantified the production of infectious virus by counting plaque-forming-units (PFU) on Vero cell monolayers, as well as on the highly-permissive U2OS human bone osteosarcoma epithelial cell line (65). No differences in virus production were noted between the four isolates when quantified on either cell type (Figure 2D, E). U2OS cells lack innate sensing of viral infection through the STING pathway (66) and can even support the growth of highly-defective HSV isolates that lack ICP0 function (65). All isolates formed large plaques on the highly-permissive U2OS cell monolayers (Figure S1), allowing us to rule out the possibility that very small foci of infection were missed during titering of the small plaque-forming isolates on Vero cells (Figure 2D, E). Finally, we compared viral protein production for these isolates, and found no differences in the expression levels of a panel of HSV-2 viral proteins at either early (6 h post infection, hpi) or late (24 hpi) time points of a single round of high MOI infection (Figure 2F, G; see legend for list of proteins). Taken together, these results suggested that large- and small-plaque-forming neonatal HSV-2 isolates did not differ significantly in viral entry, DNA replication, infectious virus production, or protein production over a single round of infection.

### Large-plaque-forming isolates exhibit enhanced cell-to-cell spread

We next assessed the ability of representative large- and small-plaque-forming isolates to spread from cell-to-cell. Vero cell monolayers were infected at low MOI (MOI = 0.001) in order to assess differences over multiple rounds of viral replication and spread throughout the cell monolayer (Figure 3A). The contribution of indirect cell-free spread was minimized by including 0.1% human serum in the media, and changing the media every 24 h to remove released virus and refresh serum levels. Over a 72-hour time course, cell monolayers were assessed to measure the extent of cell-to-cell spread, either by harvesting and titering PFU production (Figure 3B, C), or by fixing infected cell monolayers and evaluating the distribution of virus by immunofluorescence (Figure 3D, E, and Figure S2). Viral titers recovered from harvested cells were similar at 2 hpi, confirming that equivalent amounts of each virus were present following initial infection (Figure 3B). By 72 hpi, after multiple rounds of replication and spread, the large-plaque-forming isolates CNS11 and CNS03 had achieved viral titers significantly greater than those of the small-plaque-forming isolates CNS12 and SEM02 (Figure 3B, C; two-way ANOVA followed by Tukey’s multiple comparison test, p < 0.0001 at 72 h). Isolate CNS11, which was obtained directly from the CSF, produced titers statistically greater than the other three isolates (Figure 3B, C).

**Figure 3.**
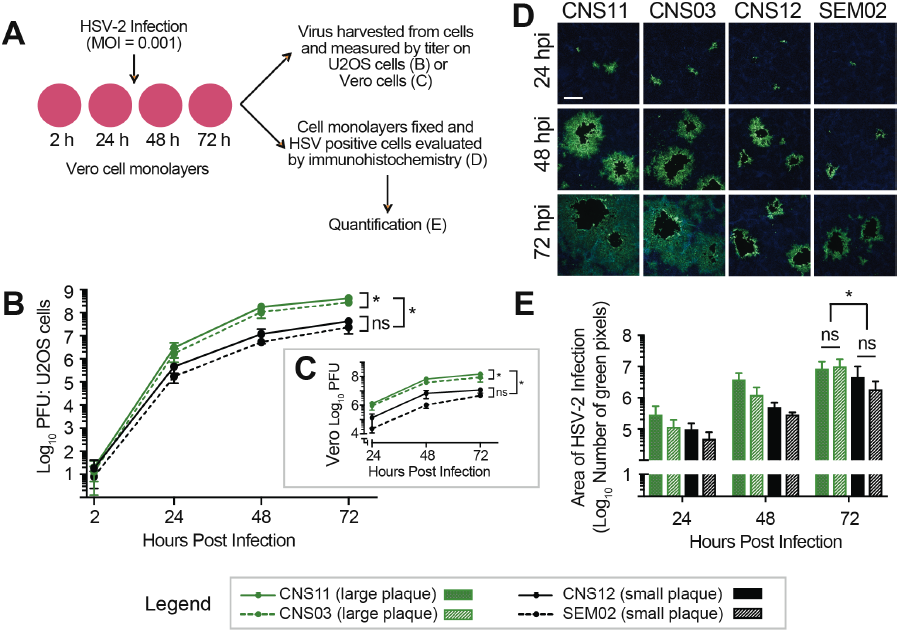
Enhanced viral cell-to-cell spread contributes to increased plaque size in culture. (**A**) The rate of viral spread in Vero cells was compared for representative neonatal isolates, including large-plaque-formers (green) and small-plaque-formers (black). Vero cell monolayers were infected at MOI = 0.001 in the presence of 0.1% human serum, which was replenished every 24 h. Samples were harvested at each time point and viral titer was evaluated by plaque formation on U2OS cells **(B)** or Vero cells **(C)**. These data represent three independent experiments. Two-way ANOVA followed by Tukey’s multiple comparison test, *p < 0.0001 at 72 h. **(D)** In parallel experiments, HSV positive cells (green) were evaluated by immunofluorescence. Cell nuclei are counterstained with DAPI (blue). Scale bar = 200 μm. Images are representative of three independent experiments. Images of the entire 10 mm coverslips were then captured and stitched to create a composite image (see Figure S2). (**E**) The total number of immunofluorescent (green) pixels was quantified for each coverslip. Two-way ANOVA followed by Tukey’s multiple comparison test, *p < 0.05 at 72 h.

We also directly evaluated cell-to-cell spread by immunostaining and quantifying the distribution of infected cells around each infectious focus. Infected Vero cell monolayers were fixed at 24, 48, and 72 hpi, and subjected to fluorescent immunocytochemistry with an antibody directed against total HSV (Figure 3D and Figure S2). The region of HSV-positive cells surrounding a single initial infection was greater for the large-plaque-forming neonatal HSV-2 isolates at 24 hpi, and increased by 48 and 72 hpi. The central cytolytic clearings seen in the monolayers infected by CNS11 or CNS03 were approximately two-fold greater than those infected by CNS12 or SEM02, reflecting the average two-fold increase in plaque size observed in methylene-blue-stained monolayers in Figure 1. However, the region of infected cells surrounding the central cytolytic clearing for CNS11 and CNS03 was dramatically larger than for CNS12 and SEM02, suggesting that large-plaque-forming neonatal HSV-2 isolates show greater spread from cell-to-cell than could have been predicted by measuring plaque size alone. To quantify this increase in the area of infected cells after a low-MOI infection, each coverslip was imaged (Figure S2) and the total number of immunofluorescent pixels was quantified (Figure 3E). By 72 hpi, the area of HSV-infected cells was statistically greater for the large-plaque-forming isolates CNS11 and CNS03, as compared to the small-plaque-forming isolates CNS12 and SEM02 (two-way ANOVA followed by Tukey’s multiple comparison test, p < 0.05 at 72 h). Together these data indicated that large-plaque-forming isolates shared an enhanced ability to spread cell-to-cell through culture, in comparison to small-plaque-forming isolates.

### Comparative genomics reveals genetic diversity in neonatal HSV-2 isolates

The differences identified in cell-to-cell spread between neonatal isolates in culture indicated the existence of intrinsic differences between these viruses. To reveal how genetic variation may contribute to viral phenotypes in culture, and ultimately to clinical disease manifestations, we sequenced the complete viral genome of all ten neonatal HSV-2 isolates. For each isolate, we sequenced purified viral nucleocapsid DNA and assembled a consensus genome, which represents the most common genotype at each nucleotide locus in the viral population. The clinical trials utilized in this study enrolled HSV-infected infants from multiple sites across the United States (8, 9, 61). Therefore, we first assessed the overall degree of relatedness between these viral genomes to understand whether any similarities in viral or geographic origin might have contributed to *in vitro* or clinical phenotype patterns. In light of the known potential for recombination in the phylogenetic history of HSV (49, 67, 53, 42), we used a graph-based network to investigate the phylogenetic relationship between these isolates. We found a similar degree of divergence among all ten neonatal HSV-2 consensus genomes (Figure 4A). We then compared these ten neonatal HSV-2 genomes to all available HSV-2 genomes in GenBank (all of which are adult; see **Table S1** for HSV-2 GenBank accessions and references) to discern any geographic or other clustering (see Table 1). The neonatal isolates did not segregate into any exclusive groupings, but instead intermingled among non-neonatal isolates, with deep branches between each isolate (Figure 4B). These findings were corroborated using an alternative phylogenetic clustering algorithm as well (Figure S3). This suggested that similarities in viral genetic origin were not responsible for determining the cellular or clinical outcomes of neonatal HSV-2 infection.

**Figure 4.**
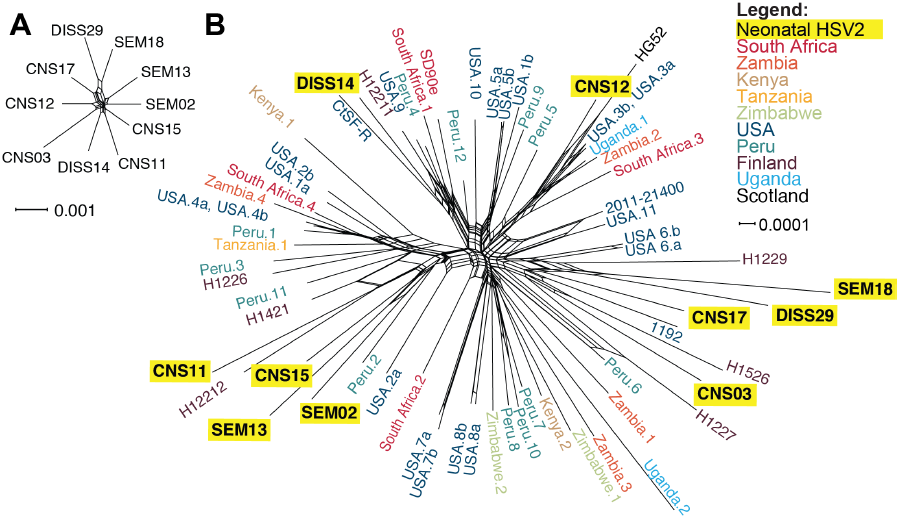
Neonatal HSV-2 genomes are genetically distinct from one another and encompass a broad range of known HSV-2 genetic diversity. A phylogenetic network constructed among neonatal HSV-2 genomes (**A**), or neonatal and adult HSV-2 genomes (**B**), reveals the wide genetic distribution of these unrelated isolates. HSV-2 genomes have been previously noted to lack geographic separation into clades (41, 42, 82). The network was created using SplitsTree4, from a MAFFT trimmed genome alignment. See Figure S3 for a comparison tree constructed using a neighbor-joining (NJ) algorithm. See **Table S1** for a complete list of accessions, geographic origins, and references for all 58 adult HSV-2 strains.

### Overall protein-coding diversity in neonatal HSV-2 isolates is similar to that observed in adult HSV-2 strains

We next asked whether overt defects in any single HSV-2 protein might be associated with clinical or *in vitro* spread phenotypes. In examining the coding potential of all ten neonatal HSV-2 isolates, we found no protein deletions or truncations encoded by any of these viral genomes. These comparisons revealed a total of 784 nucleotide differences in 71 genes (see **Table S2**), resulting in 342 non-synonymous AA differences in 65 proteins. This led to an average of 1% coding variation (range 0.0 : 2.6%). This variation was spread widely throughout the HSV-2 genome, without concentration in a particular genomic region or category of protein function (Figure 5). We found a similar level of coding diversity in this set of neonatal isolates as in a comparable set of 10 adult HSV-2 isolates (Figure 5, Table S2). We also compared these neonatal genomes to the full set of 58 annotated adult HSV-2 genomes from GenBank (listed **Table S1**). As expected from the difference in sample size, there was more overall diversity found across 58 adult HSV-2 genomes than in 10 neonatal genomes. A comparison of the dN/dS ratio in neonatal vs. adult HSV-2 genomes revealed a similar trend with a few outliers visible on each axis, e.g. UL38 and US8A had a higher dN/dS ratio in neonatal than in adult isolates (see **Table S2** and Figure S4 for full comparison). These data indicated that at the consensus level, neonatal HSV-2 isolates display substantial inter-host coding diversity spread throughout the genome, but do not possess strikingly more diversity or an excess of genetic drift as compared to adult isolates.

**Figure 5.**
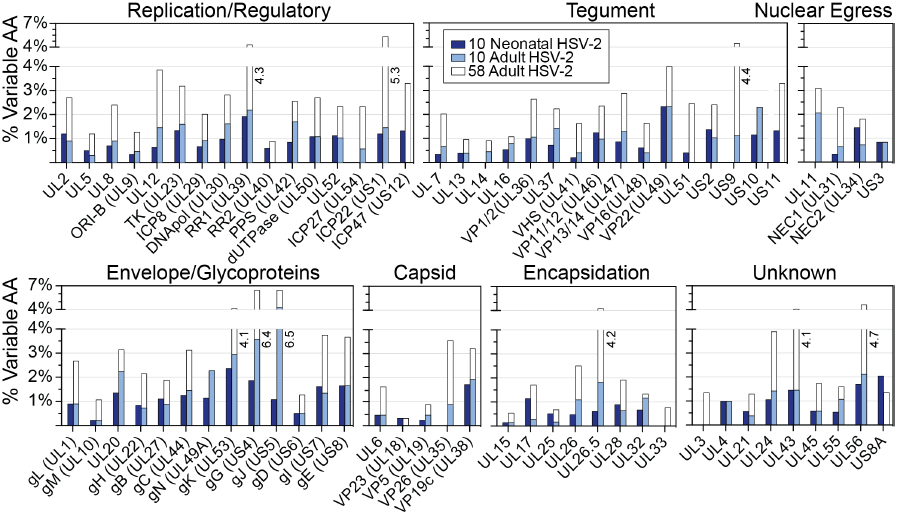
Neonatal HSV-2 proteins contain a similar degree of consensus-level amino acid (AA) differences as adult HSV-2 isolates. HSV-2 proteins are grouped by function, and the percentage of variable AAs in each protein (#AA differences divided by protein length) are plotted for the 10 neonatal isolates (dark blue), for 10 representative adult HSV-2 isolates (light blue), and for all 58 annotated adult HSV-2 genomes in GenBank (clear outlines behind light blue boxes). All adult HSV-2 genomes used for this comparison are listed in **Table S1**.

**Table 2.** Genome sequencing statistics for neonatal HSV-2 strains.

| **Clinical Isolate\*** | **Average Coverage** | **Raw Sequence Reads** | **Used for Assembly\*\*** | **% viral** | **Depth > 100** | **GenBank Accession** |
| --- | --- | --- | --- | --- | --- | --- |
| **CNS11** | 6,081X | 3.5 million | 2.8 million | 79% | 96% | MK105996 |
| **DISS14** | 6,267X | 3.8 million | 3.0 million | 77% | 97% | MK106000 |
| **CNS03** | 6,269X | 3.9 million | 3.1 million | 78% | 96% | MK105995 |
| **CNS15** | 7,486X | 6.1 million | 4.4 million | 73% | 97% | MK105998 |
| **CNS17** | 5,059X | 3.1 million | 2.3 million | 73% | 96% | MK105999 |
| **DISS29** | 2,588X | 1.4 million | 1.1 million | 78% | 99% | MK106001 |
| **CNS12** | 5,839X | 3.9 million | 2.8 million | 73% | 96% | MK105997 |
| **SEM02** | 6,455X | 4.8 million | 3.7 million | 77% | 95% | MK106002 |
| **SEM13** | 5,291X | 3.4 million | 2.4 million | 72% | 89% | MK106003 |
| **SEM18** | 7,514X | 16.9 million | 12.2 million | 72% | 98% | MK106004 |

\* Clinical isolate order based on **Figure 1**.
\*\* Numbers reflect the count of sequence read 1 of paired-end reads.

### Minor variants expand the potential coding diversity of neonatal HSV-2 isolates

We next focused our attention on differences below the consensus level in each intra-host viral population. The amino acid (AA) variations described above exist in the consensus genomes of each isolate. Since viral replication creates a population of genomes, we next assessed whether minor allelic variants existed within the viral population of any neonatal HSV-2 isolate, thereby expanding the viral genetic diversity within each host. The significant depth of coverage from deep-sequencing of each isolate allowed us to search for minor variants at every nucleotide position of each genome. We defined a minor variant as any nucleotide allele (single nucleotide polymorphism, or SNP) or insertion/deletion (INDEL) with frequency below 50%, but above 2% (a conservative limit of detection; see Methods for additional criteria). We found minor variants in the viral genome population of all 10 neonatal HSV-2 isolates, albeit to a different degree in each isolate (Figure 6A and Table S3). In total, there were 1,821 minor variants, distributed across all genomic regions (Figure S5**; Table S3**). For both SNPs and INDELs, intergenic minor variants outnumbered those in genes (genic), likely reflecting the higher selective pressures against unfavorable mutations in coding regions. The neonatal isolate DISS29 had 8-10-fold higher levels of minor variants than other neonatal isolates (Figures 6A and S5), and these variants were often present at a higher frequency or penetrance of the minor allele than observed in other neonatal isolates (Figures 6B and S5). We further examined the distribution of minor variants that occurred in genes, and found that nearly every HSV-2 protein harbored minor variants in at least one neonatal isolate (Figure 6C). Only UL3, UL11, UL35, and UL55 were completely devoid of minor variants. Three of these genes (UL3, UL11, UL35) were also devoid of AA variations at the consensus level (Figure 5). These data revealed the breadth of potential contributions of minor variants to neonatal HSV-2 biology, which could undergo selection over time or in specific niches.

**Figure 6.**
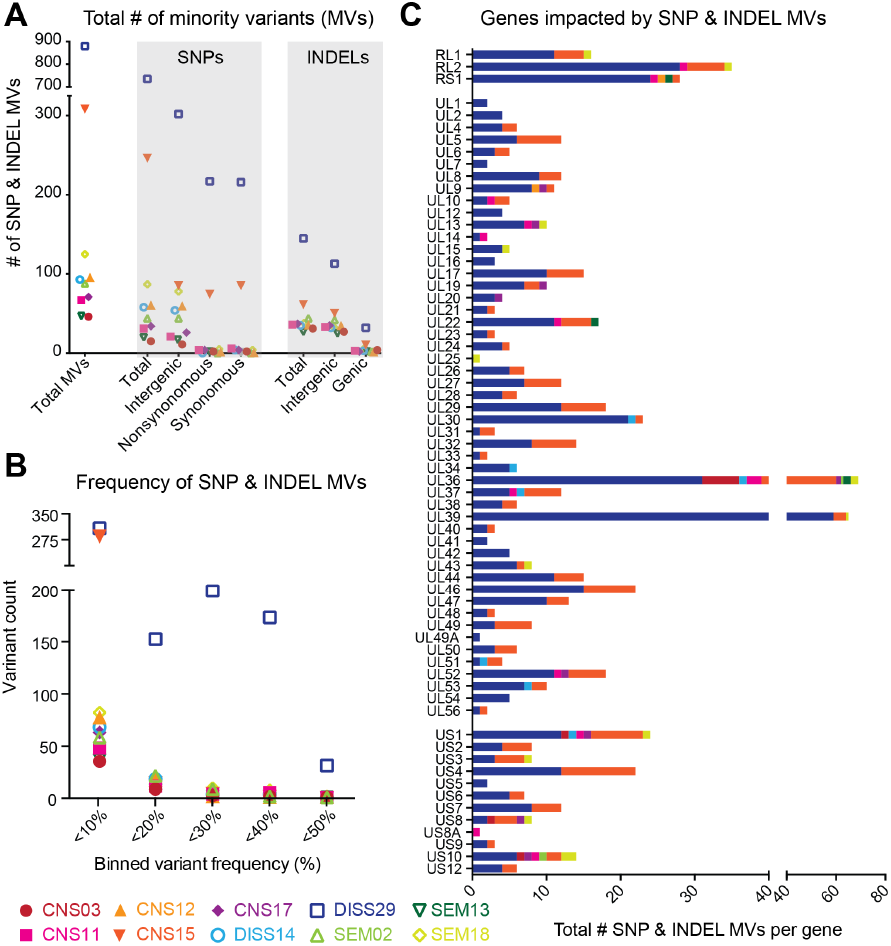
Minor variants expand the range of neonatal HSV-2 coding diversity. (**A**) Scatter plot indicates the total number of minor variants (MV; y-axis) observed in each neonatal isolate. MV are rare alleles that exist within each viral population, below the level of the consensus genome. The total number of MV on the left is separated into single-nucleotide polymorphism (SNP) vs. insertion/deletion (INDEL) variants on the right (x-axis). The genomic location of each SNP or INDEL variant is also summarized: intergenic vs. inside of genes (genic) for INDELS, and intergenic vs. nonsynonymous or synonymous SNPs inside of genes. (**B**) The frequency, or penetrance, of each minor variant was examined for each isolate. Data (x-axis) was binned in increments of 5% (e.g. 0-<5% frequency, 5-<10% frequency, and so on), and plotted according to the number of MV observed at each frequency (y-axis). SNP and INDEL variants were combined for this analysis. (**C**) Stacked histograms show the number of genic MV (x-axis) located in each HSV-2 coding sequence (gene; y-axis). SNP and INDEL variants were combined for this analysis. UL3, UL11, UL35, and UL55 lacked any minor variants and are not included in the histogram. See **Table S3** for full list of SNP and INDEL MV position and frequency data.

### Coding variations identified in neonatal HSV-2 isolates associated with CNS disease

To understand how viral genetic variants might relate to clinical disease, we assessed whether any of the consensus level coding variations identified in our group of ten neonatal HSV samples segregated with clinical disease features. A number of AA variations were shared by the CSF-derived isolates CNS11 and DISS14, which formed the largest plaques *in vitro*. These included AA variants at the level of the consensus genome in the HSV-2 genes encoding glycoprotein K (gK, UL53 gene: V323M), glycoprotein I (gI, US7 gene: R159L and P215S), UL8 (R221S), US2 (F137L), and glycoprotein G (gG, US4 gene: R338L, S442P, and E574D) (Figure 7). The gI variants exist individually in other viral isolates obtained from neonates with CNS disease (**Table S4**). Isolates collected from infants experiencing disseminated disease with CNS involvement also shared a variant in the HSV-2 UL20 gene (P129L) (Figure 7). One variant in the HSV-2 UL24 gene (V93A) was shared only by isolates from infants with SEM disease, with all isolates from infants with CNS disease containing a valine at this position (Figure 7). The sample size of these comparisons was constrained by the overall limits of neonatal HSV-2 isolate availability, and is too small to evaluate statistical significance for any of these associations. While CNS11 and DISS14 shared several variants listed in Figure 7, these viral genomes were not similar at the consensus genome level (Figure 4). These isolates also had non-shared coding differences in the same genes that contained shared variants (e.g. gK, UL8, gG). This indicates a potential for convergent evolution at these loci. Many of the coding variations identified in this dataset involve viral proteins that are known to modulate cell-to-cell spread (68–72) and/or contribute to neurovirulence in mouse models of CNS infection (70, 73–79) (Figure 7 and Table S4), however, their role in human disease has not yet been assessed.

**Figure 7.**
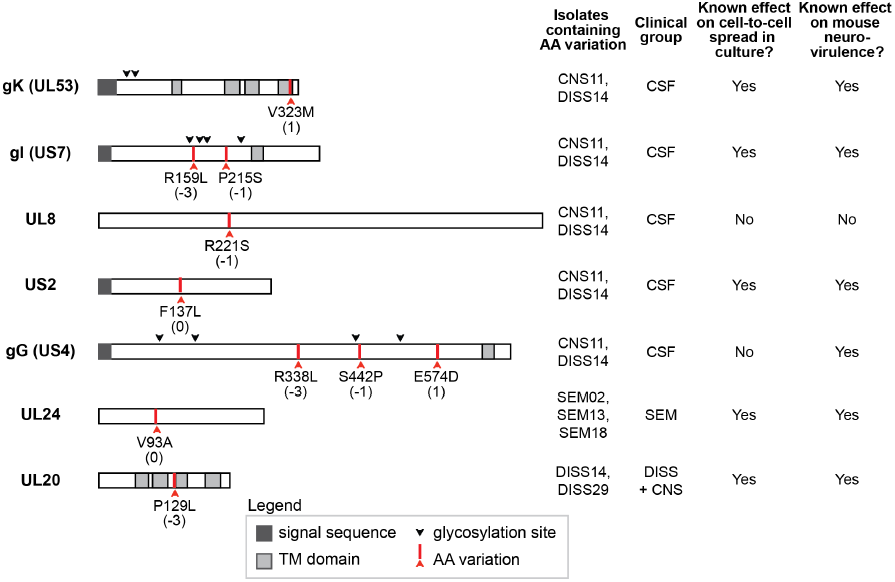
Several coding variations in neonatal HSV-2 isolates occur in proteins known to contribute to cell-to-cell spread and neurovirulence. The domain structure shown for each HSV protein is based on published literature for both HSV-1 and −2. Red arrows indicate protein-coding variations discussed in the text, with the BLOSUM80 score for each AA substitution listed in parentheses beneath the highlighted difference. More detailed information and references for each protein on the domain structure and regarding cell-to-cell spread and neurovirulence can be found in **Table S4**.

## Discussion

Host factors have not been identified to explain the >50% of neonates experiencing invasive CNS or disseminated forms of HSV infection. There is growing evidence that most herpesviruses, including HSV-1 and HSV-2, contain significant genetic variation (27, 53, 56, 80–82). The potential contributions of viral genetic variation to clinical disease in neonates therefore warrant exploration. Here, we analyzed genetic and phenotypic diversity for HSV-2 isolated from ten neonatal patients spanning two clinical studies (8, 9, 61). Although this group is small, the associations identified here provide the first insights into the potential impact of viral variability on clinical outcomes in neonatal HSV disease, and serve as a starting point for further mechanistic investigation. We found that these ten neonatal HSV-2 isolates exhibited diverse growth characteristics in culture, with larger plaque-forming isolates observed more often in infants with CNS as opposed to SEM disease. Furthermore, we established that enhanced viral spread through culture was the main contributor to this large plaque formation. Using comprehensive comparative genomics, we further demonstrated that these neonatal HSV-2 isolates contained extensive genetic diversity both within and between hosts. These data revealed several specific viral genetic variations that were associated with cases of CNS disease, in proteins known to contribute to cell-to-cell spread and/or neurovirulence in mouse models of CNS disease. Further studies are required to determine the impact of these variations on HSV-2 neurovirulence and progression to CNS disease.

Genomic comparison of these neonatal isolates revealed a wide range of genetic diversity. At the consensus genome level, which reflects the most common allele in each viral population, we found that coding differences between strains were as numerous as between previously described sets of adult HSV-2 isolates (Figure 5) (80–82). Furthermore, the specific genetic variations associated with neonatal CNS disease in our study can also be found in genital HSV-2 isolates which are not associated with CNS disease in adults. This suggests that the dramatic differences in clinical manifestations following HSV-2 infection in neonates, with significantly higher rates of invasive CNS infection, are not due to unique neonatal HSV-2 strains. However we did detect several outliers in the comparison between the rate of non-synonymous to synonymous nucleotide differences (the dN/dS ratio) of several genes in neonatal HSV-2 isolates versus those previously sequenced from adult patients (Figure S4). The genes encoding the putative virulence factor US8A (83, 84) and the capsid triplex protein VP19C (UL38) (85, 86) have a markedly higher dN/dS ratio in neonates, while that of glycoprotein C (gC; UL44) is notably lower in neonates than in adults (Figure S4). These differences in dN/dS ratio may reflect the distinct host environment of these isolates. For example viruses isolated from adult patients, often with recurrent genital infection, may have increased diversity of surface proteins such as gC due to selection for viral immune evasion (87, 88). The lower variability in gC (UL44) in neonates could result from their immunologically naïve state, and/or the shorter duration of neonatal infection prior to virus isolation (Table 1). The genes observed to have higher dN/dS ratios in neonatal isolates than in adults could result either from a loosening of selective pressures (i.e. drift) or from driving forces unique to the neonatal environment that remain to be understood. The comparison of additional isolates and ongoing characterization of under-studied proteins such as US8A (84) will help to distinguish these possibilities.

At the level of specific genetic variations in individual proteins, we detected a few fully penetrant patterns that distinguish one clinical group from another. Figure 7 highlights all of the fully-penetrant, group-specific variations that we detected in the two CSF-derived strains, two disseminated strains, and three SEM strains available for study. However other loci of potential interest exist if we consider those genetic variants found in a majority, but not all, of the neuroinvasive (CNS and DISS) strains – e.g. variations in the envelope glycoprotein gH (UL22), the viral serine/threonine kinase US3, the viral thymidine kinase (UL23), the major capsid protein UL19, and the DNA polymerase processivity factor UL42. The characterization of additional neonatal isolates will no doubt improve the clarity of these comparisons, and help to distinguish consistent patterns from those detected by chance due to the clinical limitations of the infant sample pool. Regardless of the specific viral genetic candidates under consideration, we hypothesize that viral variations impacting neonatal outcomes could either act by conferring enhanced neurovirulence, or alternatively, that they could act by limiting the rate of viral spread or degree of neuroinvasion. For instance, it is tempting to speculate that variations in glycoprotein I (US7) might be associated with enhanced viral spread, due to their observation in several large-plaque-associated viruses in this study, and prior data demonstrating the involvement of gI in both immune evasion and neuronal spread (see **Table S4**) (69, 89, 90). Conversely, the UL24 V93A variant, which was only observed in SEM-derived isolates and is relatively rare among adult HSV-2 isolates as well, could be a potential example of a spread-limiting variation. UL24 function is required to disperse nucleolin during lytic HSV-1 infection (91, 92), and mutation of UL24 has been associated with a loss of neuroinvasion in animal models (39–41). Further research will be needed to build stronger genetic associations with additional neonatal isolates, and to expand upon prior studies by testing these specific genetic variations in animal models. The application of animal models of neonatal infection will also enable the exploration of how non-genetic environmental factors such as dose and timing of viral exposure contribute to the severity and progression of disease, and how viral genetic variations and environmental factors intersect with the host genetic background.

At the level of minor variants (MV), which represent rare alleles that exist within each intra-host viral population, we found that the DISS29 harbored 8-10 fold and CNS15 harbored 3-4 fold more MV than other neonatal virus genomes (Figures 6 and S4). This could be indicative of a mixed viral population (e.g. a multi-strain infection), decreased polymerase fidelity, or the presence or absence of host selective pressure (28, 27). Diversity in viral populations has also been observed in congenital HCMV infection (30–35). However, this is the first time that evidence has been found for this level of viral population diversity with HSV-2. These minor genotypes may be selected or genetically isolated in particular niches (e.g. CSF), as observed in a comparison of VZV skin vesicles (26), or by antiviral drug selection, as was recently demonstrated in two adults with genital HSV-2 infection (93). All of the isolates sequenced in this study were collected at the initial time of diagnosis (≤19 days old), prior to acyclovir treatment and prior to the development of an HSV-specific immune response. It would therefore be compelling to examine the viral genome population from serial patient isolates over time to identify shifts in the frequency of MV due to antiviral or immune selection. Isolates from different body sites of the same patient could likewise be compared to determine whether particular genotypes are enriched in different body niches.

This comparison of viral genotype to clinical phenotype revealed associations between neonatal CNS disease and several viral protein variants that may impact neurovirulence through modulation of cell-to-cell spread. Although the sample set in this proof of concept study is small, we observed potential patterns that warrant exploration in a larger dataset. It is important to acknowledge that there is limited availability of samples from neonatal infection, both due to the rarity of these infections, and the fragility and limited body size of the infected infants. These natural circumstances lead to minimal sample collection from infected neonates. The finding that CNS-associated isolates in our study often exhibit enhanced spread between cells in culture, particularly those derived directly from the CSF, suggests that one or more of these variants could be functionally significant. Coding differences in viral proteins not known to contribute to neurovirulence were also found to be associated with neonatal CNS disease, and represent potential novel contributions to invasive infection. These promising results warrant exploration in a larger study, ideally with isolates from multiple time points and/or body sites from each infected infant. This would enable a better understanding of how overall viral genetic diversity contributes to neuroinvasion.

## Methods

### Viruses

Viruses were collected from neonates enrolled in clinical studies (8, 9, 61) by the Collaborative Antiviral Study Group (CASG) at the University of Alabama Birmingham (UAB). Samples were collected from either the cerebrospinal fluid (CSF) or skin. Enrollment in original studies was evenly split between males and females, and included both black and white patients. Clinical morbidity score was determined at 12 months of life as previously defined (94, 12). Initial collection of samples, use of samples in this study, and use of de-identified clinical information was approved by the UAB Institutional Review Board. See Supplemental Information for additional details of this and subsequent methods.

### Cell culture

Human lung fibroblast MRC-5 cells (ATCC®, CCL-171), African green monkey kidney Vero cells (ATCC®, CCL-81), and human epithelial bone osteosarcoma U2OS cells (ATCC®, HTB-96) were cultured under standard conditions. Cell lines were authenticated by ATCC prior to purchase, and were confirmed to be mycoplasma free throughout experiments by periodic testing (LookOut Mycoplasma, Sigma).

### Virus culture

Viruses were cultured at the time of diagnosis and snap frozen after 1 passage. Each viral isolate was then passaged 3 times on MRC-5 cells at a MOI of 0.01, with harvest at the time of complete cytopathic effect (between 50-70 h). Viral stocks were titered on either Vero cells (100 h) or U2OS cells (48 h) under a methylcellulose overlay. Plaque size and morphology did not change for any viral isolates over the course of virus stock expansion.

### Plaque measurements

Plaques were stained with 0.5% methylene blue. Serial 4X brightfield images were collected on an EVOS FL Auto Imaging System and stitched by EVOS software (University of Pennsylvania Cell and Developmental Biology Microscopy Core). Plaque area was measured using ImageJ software.

### Genome copy number estimation by quantitative PCR for UL27

DNA was extracted and quantitative PCR was performed with primers specific to the viral glycoprotein B gene (gB; UL27) (43). Absolute quantification was calculated based on a standard curve of HSV-1 strain F nucleocapsid DNA (59).

### Viral entry assay

Monolayers of Vero cells were cooled to 4°C for 30 minutes prior to infection with 100 PFU of each viral isolate. After 1 h of viral incubation at 4°C, unbound virus was removed by washing and cells were moved to 37°C. At 0, 10, 20, 30, 45, or 60 minutes, a low-pH citrate buffer was applied to inactivate extracellular virus. For each condition, parallel infections were performed without the addition of citrate solution (control). Cell monolayers were washed and allowed to form plaques under methylcellulose for 100 h. Viral entry was quantified as the fraction of plaques formed following citrate buffer application, where 100% is the number of plaques formed on a monolayer not treated with citrate buffer (control).

### Single-step and multi-step growth curves

Growth curves of HSV infection were performed in Vero cells. At 2 hpi, 0.1% human serum was added to reduce cell-free spread of virus. Single-step growth curves were performed at MOI = 5, and multi-step growth curves at MOI = 0.001, as defined by titering viral stocks on U2OS cells. Every 24 h, the supernatant was removed and media containing 0.1% human serum reapplied.

### Immunocytochemistry

Immunocytochemistry was performed as previously described (95). Infection was detected with rabbit anti-HSV primary antibodies (Agilent Dako, B0114) and fluorophore-conjugated anti-rabbit secondary antibodies (Invitrogen, A-11008). Cell nuclei were counterstained with DAPI. 5X images were collected with a Leica DM6000 wide field microscope equipped with a Photometrics HQ2 high resolution monochrome CCD camera, and processed with LAS AF software (UPenn Cell and Developmental Microscopy Core). 10X images were collected on an EVOS FL Auto Imaging System and stitched using EVOS software (UPenn Cell and Developmental Microscopy Core). Exposure and gain were optimized within each experiment for one virus at the 72 h time point and applied identically to each image within that experiment. Subsequent image processing (ImageJ) was applied equally to all images in a given experiment.

### Immunoblotting

Immunoblotting was performed as previously described (95). Equal amounts of whole cell lysate were separated by SDS-PAGE. Membranes were immunoblotted with antibodies raised against total HSV (Agilent Dako, B0114); glycoproteins (g)C, gD, gE, gH, and VP5 (all gifts from Gary Cohen); ICP8 (gift from David Knipe); and GAPDH (GeneTex, GTX100118).

### Viral DNA isolation and Illumina sequencing

Viral nucleocapsid DNA for genome sequencing was prepared by infecting MRC-5 cells at an MOI ≥ 5 as previously described (96, 97). Viral nucleocapsid gDNA was sheared using a Covaris M220 sonicator/disruptor under the following conditions: 60s duration, peak power 50, 10% duty cycle, at 4°C. Barcoded sequencing libraries were prepared using the Illumina TruSeq low-throughput protocol according to manufacturer’s specifications and as previously described (60, 98). The quality of sequencing libraries was evaluated by Qubit (Invitrogen, CA), Bioanalyzer (Agilent), and qPCR (KAPA Biosystems). Paired-end sequencing (2 × 300 bp length) was performed on an Illumina MiSeq, according to manufacturer’s recommendations (17pM input).

### *De novo* genome assembly

A consensus genome was assembled for each viral isolate using a previously described Viral Genome Assembly (VirGA) bioinformatics workflow (60). Annotation of new genome sequences was guided by the HSV-2 reference genome (strain HG52; GenBank NC_001798) based on sequence homology (99).

### Comparative genomics and phylogenetic analysis

The 10 neonatal HSV-2 genomes were aligned with all annotated HSV-2 genomes available in GenBank (see **Table S1** for full list of 58 genomes; all derived from adults) using MAFFT (100). The genome-wide alignment used a trimmed genome format (lacking the terminal repeats) to avoid giving undue weight to these duplicated sequences. The MAFFT alignment was used to generate a NeighborNet phylogenetic network in SplitsTree with Uncorrected P distances (49, 101, 102), as well as a Neighbor-Joining tree (Jukes-Cantor, 1000 bootstraps) in MEGA Omega for Windows (103). A diverse subset of ten adult HSV-2 isolates was selected for protein-level comparisons with the ten neonatal isolates (indicated in **Table S1** with an asterisk). ClustalW2 was used to construct pairwise nucleotide alignments between whole genomes, and pairwise amino acid alignments for each gene and protein (104). Pan-HSV2 comparisons excluded three viral proteins for which sequences are not fully determined in most published strains, likely due to the high G + C-content and numerous tandem repeats in these regions: ICP34.5 (RL1 – annotated/complete in only 9 genomes from **Table S1**), ICP0 (RL2 – annotated/complete in only 8 genomes from **Table S1**), and ICP4 (RS1 – annotated/complete in only 6 genomes from **Table S1**) (80–82). Custom Python scripts were used on these alignments to identify nucleotide and AA differences between samples.

### Minor variant detection & quantification

Minor variants (MV) were detected using VarScan v2.2.11 (mpileup2snp and mpileup2indel commands) (105), using the following parameters to differentiate true MV from technical artifacts (26): minimum allele frequency ≥ 0.02 (2%); base call quality ≥ 20; read depth ≥ 100; independent reads supporting minor allele ≥5. MV with directional strand bias ≥ 90% were excluded. The genomic location and potential impact of each MV was assessed using SnpEff and SnpSift (106, 107). We use the conservative cutoff of 2% for detection of minor variants both to avoid false positive signals and to provide data that is comparable to recent examples in the genomic analysis of herpesviruses in clinical samples (108, 28, 109).

## Acknowledgements

We thank Penny Jester and the UAB clinical support team for their assistance in retrieving archival records. The UPenn Cell and Developmental Biology Microscopy Core provided imaging assistance, and Elise M. Peaurori provided technical assistance during early stages of this project. We thank members of the Szpara and Weitzman labs for helpful discussions. This research was supported by NIAID grant 1R21AI140443 (MLS and MDW), with additional support from the following: startup funds from the Pennsylvania State University (MLS), a CURE grant from the Pennsylvania Department of Health (MLS), a grant from the National Institutes of Health (NS082240 to MDW), and the NICHD Pediatric Scientist Development grant K12-HD000850 (LNA). The published NIAID CASG clinical studies of neonatal HSV infection were funded by a contract (N01-AI-62554) with the NIAID Development and Applications Branch and by grants from the General Clinical Research Center Program (RR-032) and the state of Alabama.

## Author Contributions

Writing – Original Draft: LNA, CB, AND, and MLS; Writing – Review & Editing: LNA, MDW, and MLS; Conceptualization: LNA and MLS; Investigation: LNA, CB, AND, DWR, UP; Resources: DWK, MNP, and RJW; Funding Acquisition: MLS, RJW, and MDW.

## Supplemental Information

### Virus source

Viruses analyzed were collected from neonates with HSV-2 infection, enrolled in three published clinical studies (1–3) by the Collaborative Antiviral Study Group (CASG) at the University of Alabama Birmingham (UAB). These HSV isolates originated from patients with SEM, CNS, or disseminated (DISS) disease with CNS involvement (Table 1). Samples were collected at the time of diagnosis, prior to initiation of acyclovir therapy. CNS11 and DISS14 were isolated from the cerebrospinal fluid (CSF) and all other viruses were isolated from the skin. CSF was collected from all infants at the time of diagnosis. However, for all of the viruses isolated from the skin of infants with CNS disease (CNS03, CNS12, CNS15, CNS17, and DISS29), the CSF collected at the time of diagnosis was PCR positive but culture negative for HSV. Viral isolates are associated with de-identified clinical information including age, sex, race, and clinical morbidity scores, as approved by the UAB Institutional Review Board (IRB). Enrollment in the original studies was evenly split between males and females, and included both black and white patients. Clinical morbidity score was determined by the CASG after 12 months of life as (1) normal; (2) mild impairment which includes ocular sequelae (keratoconjunctivitis), speech delay, or mild motor delay; (3) moderate impairment which includes hemiparesis, persistent seizure disorder, or developmental delay of less than or equal to 3 months adjusted developmental age; (4) severe impairment which includes microcephaly, spastic quadriplegia, chorioretinitis or blindness, or a serious developmental delay of least 3 months according to the Denver Developmental Assessment Scale (1, 2). These assessments were performed at each collaborating site. Initial collection of samples was approved by the UAB IRB. Use of these samples in this study was approved by the UAB IRB.

### Cell culture

Human lung fibroblast MRC-5 cells (ATCC®, CCL-171) were cultured in minimum essential medium Eagle (MEME, Sigma-Aldrich; M5650) supplemented with 10% fetal bovine serum (FBS, Hyclone; SH30071.03), 2mM L-glutamine (Gibco, A2916801) and 1X penicillin-streptomycin (Gibco; 15140122). African green monkey kidney Vero cells (ATCC®, CCL-81) and human epithelial bone osteosarcoma U2OS cells (ATCC®, HTB-96) were cultured in Dulbecco’s modified Eagle’s medium with high glucose (DMEM, Hyclone; SH30081.02) supplemented with 10% FBS, 2mM L-glutamine, and 1X penicillin-streptomycin. Cell lines were authenticated by ATCC prior to purchase, and were confirmed to be mycoplasma free throughout experiments by periodic testing (LookOut Mycoplasma, Sigma).

### Virus culture

Initial viral isolates (passage 1-2) were obtained from CASG samples stored at the UAB. Each isolate had been cultured at the time of diagnosis, aliquoted after one passage, and snap frozen. To conduct viral genome sequencing and phenotypic assays, each stock was expanded over three serial passages by infecting monolayers of human MRC-5 cells at a MOI of 0.01. Each infection was allowed to progress to complete cytopathic effect (CPE) before harvest (between 50-70 hours), with increasing cell volume at each passage to produce sufficient virions for subsequent phenotypic and genomic comparisons. Viral stocks were titered on monolayers of either Vero cells or U2OS cells, for 100 or 48 hours respectively, to allow plaques to develop. Plaque formation was facilitated by limiting viral diffusion with a methylcellulose overlay. Plaque size and morphology were monitored carefully and did not change for any viral isolates over the course of virus stock expansion.

### Plaque measurements

After appropriate incubation plaques were stained with 0.5% methylene blue and allowed to dry. Serial 4X brightfield images were collected on an EVOS FL Auto Imaging System and stitched by EVOS software to create an image of the entire well (University of Pennsylvania Cell and Developmental Biology Microscopy Core). No processing was performed. The area of 100 plaques was measured for each viral isolate using ImageJ software.

### Genome copy number estimation by quantitative PCR for UL27

DNA was extracted using a PureLink genomic DNA mini kit (ThermoFisher Scientific). Viral genome copy number was determined using an established assay based on real-time PCR using primers and dual-fluorescent probe specific to viral glycoprotein B gene (gB; UL27). Samples were assayed alongside a standard curve of HSV-1 strain F nucleocapsid DNA (3), on a ViiA 7 Real-Time PCR System.

### Viral entry assay

Monolayers of Vero cells were cooled to 4°C for 30 minutes prior to infection and 100 PFU of each viral isolate was applied to cell monolayers at 4°C for 1 hour to allow virus binding, after which unbound virus was washed from the cells. Cells were then moved to 37°C to allow virus entry. At 0, 10, 20, 30, 45, or 60 minutes, a low-pH citrate buffer was applied to infected cells to inactivate virus that had not penetrated the cellular membrane. At each time point and for each virus, parallel infections were performed without the addition of citrate solution. These served as controls to determine the maximum number of plaques formed. Cell monolayers were washed and overlaid with methylcellulose. Plaques were scored after 100 hours of incubation. Viral entry was quantified as the fraction of plaques formed following citrate buffer application, where 100% is the number of plaques formed on a monolayer not treated with citrate buffer (control).

### Single-step and multi-step growth curves

Virus diluted in basic growth media containing 2% FBS was applied to near confluent monolayers of Vero cells and allowed to adsorb for 1 hour. Virus was removed and cells were incubated at 37C for the duration of infection. At 2 hpi, media containing 0.1% human serum was added to reduce the contribution of cell-free spread of virus. Single-step growth curves were performed at MOI=5 as defined by titering viral stocks on U2OS cells, and monolayers harvested at 2, 6, 12, and 24 hpi. Multi-step growth curves were performed at MOI=0.001 as defined by titering viral stocks on U2OS cells, and monolayers were harvested at 2, 24, 48, and 72 hpi. Every 24 hours, the supernatant was removed and media containing 0.1% human serum reapplied. At the conclusion of each infection, cells were washed two times with PBS and collected by scraping into an equal volume of media.

### Immunocytochemistry

Multi-step growth curves were performed as described above in Vero cell monolayers grown on glass coverslips. To terminate each infection, coverslips were washed with PBS and fixed in 4% paraformaldehyde for 15 minutes at room temperature. Cells were permeabilized with 0.5% Triton-X, blocked in 3% BSA, and incubated with polyclonal rabbit antibodies raised against total HSV (Agilent Dako, B0114). After washing, cells were incubated with fluorophore-conjugated anti-Rabbit secondary antibodies (Invitrogen, A-11008) to mark HSV infected cells, and counterstained with DAPI to mark cell nuclei. Coverslips were initially visualized with a Leica DM6000 wide field microscope (UPenn Cell and Developmental Microscopy Core). The 5X images were collected on a Photometrics HQ2 high resolution monochrome CCD camera, and processed with LAS AF software. The 10X images were collected on a EVOS FL Auto Imaging System (UPenn Cell and Developmental Microscopy Core) and stitched using EVOS software. Exposure and gain were optimized within each experiment for one virus at the 72-hour time point and applied identically to each image within that experiment. Any subsequent image processing (ImageJ) was applied equally to all images in a given experiment.

### Immunoblotting

Whole cell lysates were prepared with 1X LDS sample buffer (NuPage) and equal amounts of lysate were separated by SDS-PAGE. Membranes were immunoblotted with polyclonal rabbit antibodies raised against total HSV (Agilent Dako, B0114); glycoproteins (g)C, gD, gE, gH, and VP5 (all gifts from Gary Cohen); ICP8 (gift from David Knipe); and GAPDH (GeneTex, GTX100118). Proteins were visualized with Pierce ECL Western Blotting Substrate (ThermoFisher Scientific) and detected using a G:Box imaging system (Syngene).

### Viral DNA isolation and Illumina sequencing

Viral nucleocapsid DNA for genome sequencing was prepared by infecting MRC-5 cells at an MOI ≥5 as previously described (4, 5). Viral nucleocapsid gDNA was sheared using a Covaris M220 sonicator/disruptor under the following conditions: 60s duration, peak power 50, 10% duty cycle, at 4°C. Barcoded sequencing libraries were prepared using the Illumina TruSeq low-throughput protocol according to manufacturer’s specifications and as previously described (6, 7). The quality of sequencing libraries was evaluated by Qubit (Invitrogen, CA), Bioanalyzer (Agilent), and qPCR (KAPA Biosystems). Paired-end sequencing (2 × 300bp length) was performed on an Illumina MiSeq, according to manufacturer’s recommendations (17pM input).

### *De novo* genome assembly

A consensus genome was assembled for each viral isolate using a previously described Viral Genome Assembly (VirGA) bioinformatics workflow (6). VirGA begins by quality-filtering the MiSeq sequence reads and removing sequences that match the host (human) genome. Thereafter VirGA uses a combination of SSAKE *de novo* assemblies run with differing parameters, which are then combined into a single draft genome using Celera and GapFiller (8–10). After quality-checking and iterative improvement of the genome assembly, annotations were transferred from the HSV-2 reference genome (strain HG52; GenBank NC_001798) to each new genome based on sequence homology (11).

### Comparative genomics and phylogenetic analysis

The genomes of all 10 neonatal HSV-2 isolates were combined with all annotated HSV-2 genomes available in GenBank (see Table S1 for full list) and aligned using MAFFT (12). All previously published HSV-2 genomes were derived from infected adults. The genome-wide alignment used a trimmed genome format (lacking the terminal repeats) to avoid giving undue weight to these duplicated sequences. The MAFFT alignment was used to generate a NeighborNet phylogenetic network in SplitsTree with Uncorrected P distances (13–15). A diverse subset of ten adult HSV-2 isolates was selected for protein-level comparisons with the ten neonatal isolates; these are indicated in **Table S1** with an asterisk. GenBank genomes that lacked open reading frame (ORF) and protein annotations were excluded from all comparisons (16, 17). ClustalW2 was used to construct pairwise nucleotide alignments between whole genomes and pairwise amino acid alignments for each gene and protein (18). Pan-HSV-2 comparisons excluded three viral proteins for which sequences are not fully determined in most published strains, likely due to the high G+C-content and numerous tandem repeats in these regions: ICP34.5 (RL1 – annotated/complete in only 9 genomes from Table S1) and ICP4 (RS1 – annotated/complete in only 6 genomes from Table S1), ICP0 (RL2 – annotated/complete in only 8 genomes from Table S1) (16, 19, 20). Custom Python scripts were used on these alignments to identify nucleotide and AA differences between samples.

## Supplemental Figures & Legends

**Figure S1.**
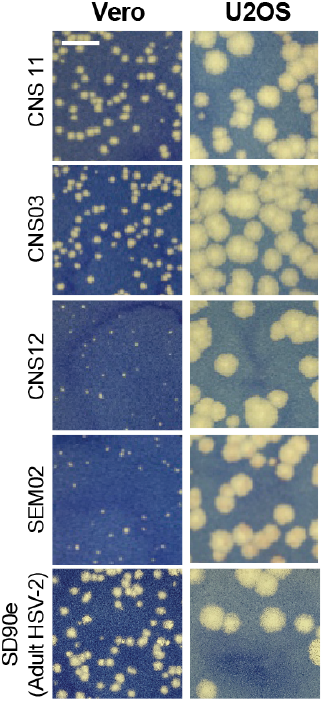
U2OS cells support large plaque formation by all isolates. Representative plaques are shown after neonatal viruses were allowed to incubate for 100 hours on Vero or U2OS cells. All neonatal isolates were capable of forming large plaques on U2OS cells, which lack innate sensing of viral infection through the STING pathway (21).

**Figure S2.**
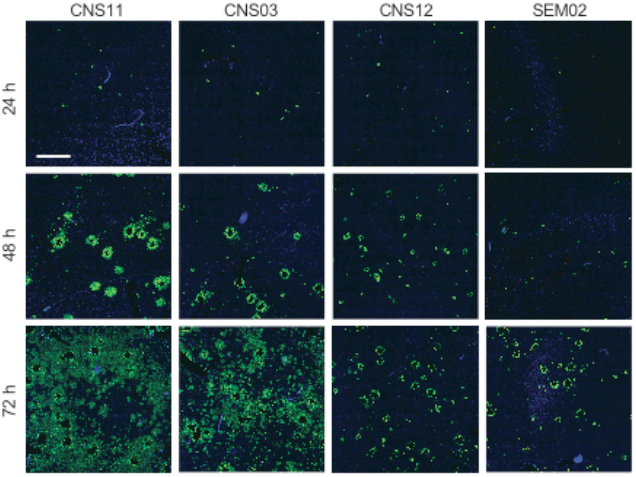
Cell-to-cell spread is enhanced in certain neonatal HSV-2 isolates. Confluent Vero cell monolayers were infected at MOI = 0.001 for the time points indicated, in the presence of 0.1% human serum. HSV positive cells (green) were evaluated at each time point by immunofluorescence. Cell nuclei are counterstained with DAPI (blue). Serial 10X images were obtained on an EVOS FL Auto Imaging System and stitched together to create an image of the entire experimental coverslip. Scale bar = 1 mm. These images were quantified in Figure 3E.

**Figure S3.**
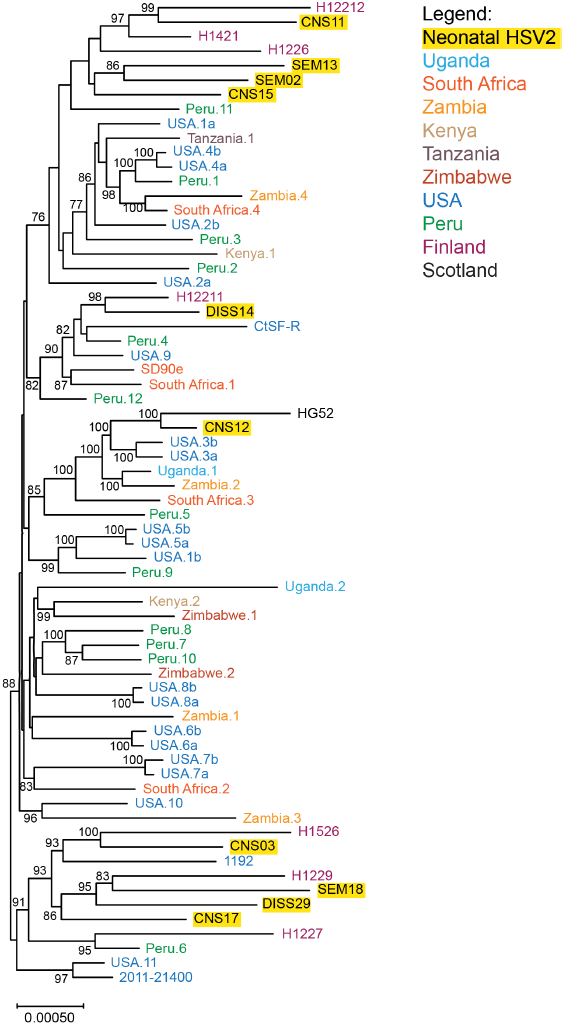
Phylogenetic clustering demonstrates that neonatal HSV-2 genomes are genetically distinct from one another and intermingle with the previously known range of HSV-2 genetic diversity. A Neighbor-Joining (NJ) tree network constructed among ten neonatal and 58 adult HSV-2 genomes, revealed the wide genetic distribution of the neonatal isolates. The NJ tree (Jukes-Cantor, 1000 bootstraps) was created in MEGA from a MAFFT trimmed genome alignment. Bootstrap values ≥ 70 are shown here. See Figure 4 for a network graph comparison to this tree.

**Figure S4.**
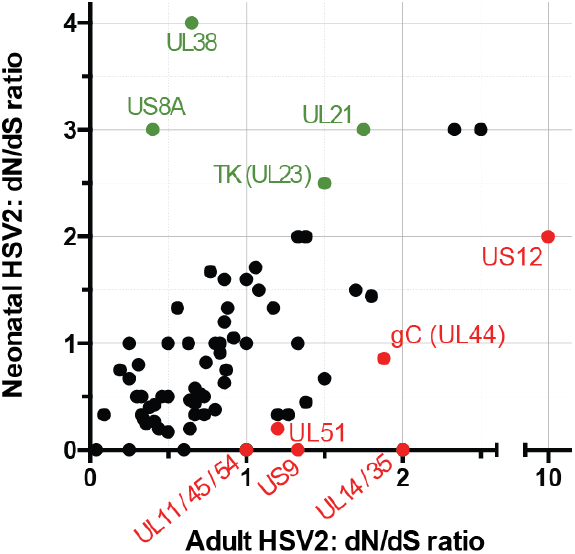
Ratio of non-synonymous to synonymous coding variations in neonatal HSV-2 versus adult HSV-2 strains. The ratio of non-synonymous (dN) to synonymous (dS) coding variations were plotted for each HSV-2 protein. The x-axis value represents the average dN/dS ratio for each protein in 58 adult HSV-2 strains, while the y-axis value represents the average dN/dS ratio in 10 neonatal isolates. Proteins with a difference in average dN/dS ratios ≥ 1 in neonatal vs. adult HSV-2 genomes are labeled (green indicates a higher average dN/dS in neonatal HSV-2 genomes; red indicates a higher dN/dS in adult HSV-2 genomes). The average dN/dS ratios for all proteins are listed in **Table S2**.

**Figure S5.**
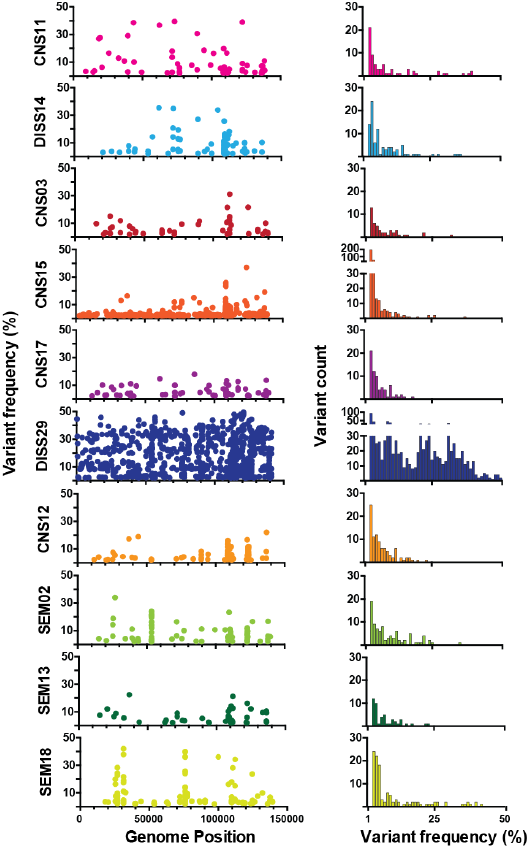
Genome-wide distribution and frequency of minor variants in neonatal HSV-2 isolates. For each neonatal HSV-2 isolate, the graph on the left plots spatial location in the genome (x-axis) against the frequency at which each minor variant was observed. The plot on the right summarizes the number of minor variants (y-axis height) in binned increments of 1% (x-axis). The data reveal the distinctly different distribution of minor variants in DISS29, and to a lesser extent CNS15, than in other isolates. Color code matches that used in Figure 6. See **Table S3** for full list of SNP and INDEL MV position and frequency data.

**Supplemental Table S1.** Adult HSV-2 genomes (58 total) used for comparative genomic analyses.

| **Virus Isolate** | **Country (with location, if available)** | **GenBank Accession #** | **References** |
| --- | --- | --- | --- |
| H1227* | Finland | KY922721 | In process |
| H1229* | Finland | KY922722 | In process |
| H12212* | Finland | KY922726 | In process |
| Uganda.2* | Uganda | KX574906 | (1) |
| Peru.8* | Peru | KX574874 | (1) |
| Zambia.4* | Zambia | KX574883 | (1) |
| HG52* | Scotland | NC_001798 | (2) |
| CtSF-R* | USA | KP334093 | (3) |
| 1192* | Wisconsin, USA | KP334095 | (3) |
| SD90e* | Carletonville, South Africa | KF781518 | (4, 5) |
| Peru.1 | Peru | KX574860 | (1, 6) |
| Peru.12 | Peru | KX574861 | (1, 6) |
| Zambia.3 | Zambia | KX574862 | (1, 6) |
| Zimbabwe.2 | Zimbabwe | KX574863 | (1, 6) |
| Zambia.2 | Zambia | KX574864 | (1, 6) |
| Peru.9 | Peru | KX574865 | (1, 6) |
| Peru.11 | Peru | KX574866 | (1, 6) |
| Peru.10 | Peru | KX574867 | (1, 6) |
| USA.8a | USA | KX574868 | (1, 6) |
| USA.8b | USA | KX574869 | (1, 6) |
| USA.1b | USA | KX574870 | (1, 6) |
| Zimbabwe.1 | Zimbabwe | KX574871 | (1, 6) |
| Peru.7 | Peru | KX574872 | (1, 6) |
| Peru.2 | Peru | KX574873 | (1, 6) |
| Peru.6 | Peru | KX574875 | (1, 6) |
| Peru.4 | Peru | KX574876 | (1, 6) |
| Peru.3 | Peru | KX574877 | (1, 6) |
| Peru.5 | Peru | KX574878 | (1, 6) |
| South Africa.2 | South Africa | KX574879 | (1, 6) |
| South Africa.3 | South Africa | KX574880 | (1, 6) |
| South Africa.4 | South Africa | KX574881 | (1, 6) |
| USA.3a | USA | KX574882 | (1, 6) |
| USA.3b | USA | KX574884 | (1, 6) |
| USA.2b | USA | KX574885 | (1, 6) |
| USA.10 | USA | KX574886 | (1, 6) |
| USA.4a | USA | KX574887 | (1, 6) |
| USA.7a | USA | KX574888 | (1, 6) |
| USA.6a | USA | KX574889 | (1, 6) |
| USA.6b | USA | KX574890 | (1, 6) |
| USA.5a | USA | KX574891 | (1, 6) |
| South Africa.1 | South Africa | KX574892 | (1, 6) |
| Kenya.2 | Kenya | KX574893 | (1, 6) |
| Kenya.1 | Kenya | KX574894 | (1, 6) |
| USA.7b | USA | KX574895 | (1, 6) |
| USA.5b | USA | KX574896 | (1, 6) |
| USA.11 | USA | KX574897 | (1, 6) |
| USA.4b | USA | KX574898 | (1, 6) |
| Uganda.1 | Uganda | KX574899 | (1, 6) |
| USA.9 | USA | KX574901 | (1, 6) |
| Tanzania.1 | Tanzania | KX574902 | (1, 6) |
| USA.1a | USA | KX574903 | (1, 6) |
| USA.2a | USA | KX574904 | (1, 6) |
| Zambia.1 | Zambia | KX574905 | (1, 6) |
| 2011-21400 | USA | KX574908 | (1, 6) |
| H12211 | Finland | KY922725 | In process |
| H1226 | Finland | KY922720 | In process |
| H1421 | Finland | KY922723 | In process |
| H1526 | Finland | KY922724 | In process |

\*Indicates 10 adult HSV-2 strains used for AA comparisons in Figure 5.

---

**Supplemental Table S2.**
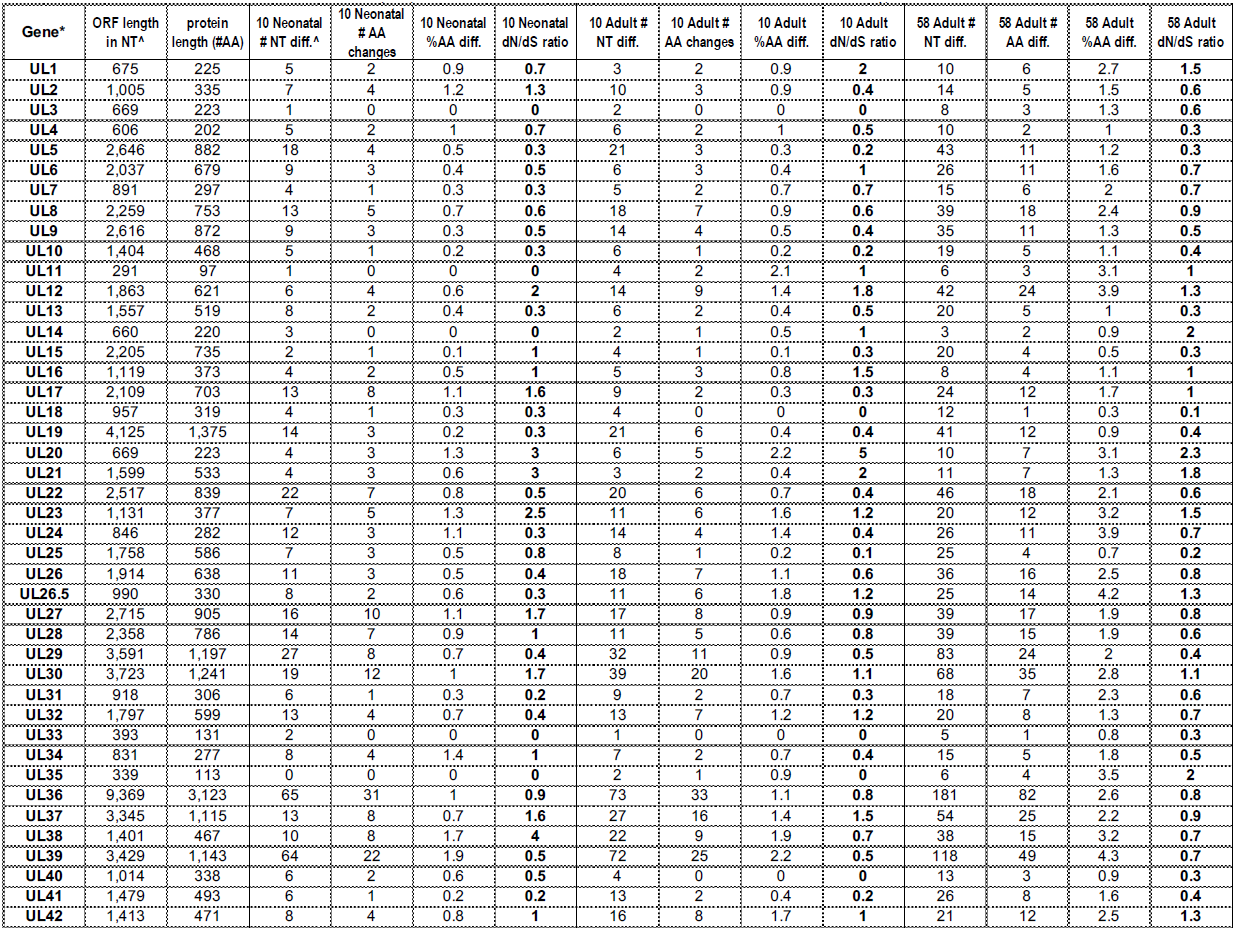
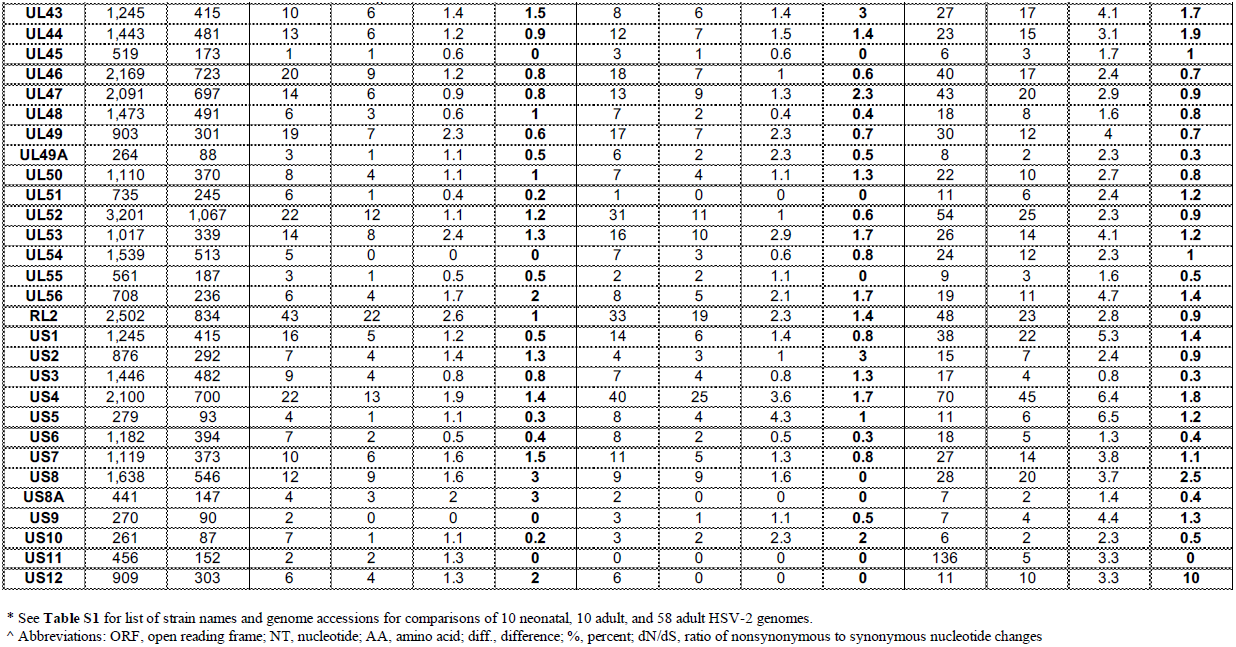
Number of nucleotide and amino acid (AA) differences observed in each set of neonatal or adult HSV-2 genomes.

| Gene* | ORF length<br>in NT<sup>A</sup> | protein<br>length (#AA) | 10 Neonatal<br># NT diff.<sup>A</sup> | 10 Neonatal<br># AA<br>changes | 10 Neonatal<br>%AA diff. | 10 Neonatal<br>dN/dS ratio | 10 Adult #<br>NT diff. | 10 Adult #<br>AA changes | 10 Adult<br>%AA diff. | 10 Adult<br>dN/dS ratio | 58 Adult #<br>NT diff. | 58 Adult #<br>AA diff. | 58 Adult<br>%AA diff. | 58 Adult<br>dN/dS ratio |
| --- | --- | --- | --- | --- | --- | --- | --- | --- | --- | --- | --- | --- | --- | --- |
| UL1 | 675 | 225 | 5 | 2 | 0.9 | 0.7 | 3 | 2 | 0.9 | 2 | 10 | 6 | 2.7 | 1.5 |
| UL2 | 1,005 | 335 | 7 | 4 | 1.2 | 1.3 | 10 | 3 | 0.9 | 0.4 | 14 | 5 | 1.5 | 0.6 |
| UL3 | 669 | 223 | 1 | 0 | 0 | 0 | 2 | 0 | 0 | 0 | 8 | 3 | 1.3 | 0.6 |
| UL4 | 606 | 202 | 5 | 2 | 1 | 0.7 | 6 | 2 | 1 | 0.5 | 10 | 2 | 1 | 0.3 |
| UL5 | 2,646 | 882 | 18 | 4 | 0.5 | 0.3 | 21 | 3 | 0.3 | 0.2 | 43 | 11 | 1.2 | 0.3 |
| UL6 | 2,037 | 679 | 9 | 3 | 0.4 | 0.5 | 6 | 3 | 0.4 | 1 | 26 | 11 | 1.6 | 0.7 |
| UL7 | 891 | 297 | 4 | 1 | 0.3 | 0.3 | 5 | 2 | 0.7 | 0.7 | 15 | 6 | 2 | 0.7 |
| UL8 | 2,259 | 753 | 13 | 5 | 0.7 | 0.6 | 18 | 7 | 0.9 | 0.6 | 39 | 18 | 2.4 | 0.9 |
| UL9 | 2,616 | 872 | 9 | 3 | 0.3 | 0.5 | 14 | 4 | 0.5 | 0.4 | 35 | 11 | 1.3 | 0.5 |
| UL10 | 1,404 | 468 | 5 | 1 | 0.2 | 0.3 | 6 | 1 | 0.2 | 0.2 | 19 | 5 | 1.1 | 0.4 |
| UL11 | 291 | 97 | 1 | 0 | 0 | 0 | 4 | 2 | 2.1 | 1 | 6 | 3 | 3.1 | 1 |
| UL12 | 1,863 | 621 | 6 | 4 | 0.6 | 2 | 14 | 9 | 1.4 | 1.8 | 42 | 24 | 3.9 | 1.3 |
| UL13 | 1,557 | 519 | 8 | 2 | 0.4 | 0.3 | 6 | 2 | 0.4 | 0.5 | 20 | 5 | 1 | 0.3 |
| UL14 | 660 | 220 | 3 | 0 | 0 | 0 | 2 | 1 | 0.5 | 1 | 3 | 2 | 0.9 | 2 |
| UL15 | 2,205 | 735 | 2 | 1 | 0.1 | 1 | 4 | 1 | 0.1 | 0.3 | 20 | 4 | 0.5 | 0.3 |
| UL16 | 1,119 | 373 | 4 | 2 | 0.5 | 1 | 5 | 3 | 0.8 | 1.5 | 8 | 4 | 1.1 | 1 |
| UL17 | 2,109 | 703 | 13 | 8 | 1.1 | 1.6 | 9 | 2 | 0.3 | 0.3 | 24 | 12 | 1.7 | 1 |
| UL18 | 957 | 319 | 4 | 1 | 0.3 | 0.3 | 4 | 0 | 0 | 0 | 12 | 1 | 0.3 | 0.1 |
| UL19 | 4,125 | 1,375 | 14 | 3 | 0.2 | 0.3 | 21 | 6 | 0.4 | 0.4 | 41 | 12 | 0.9 | 0.4 |
| UL20 | 669 | 223 | 4 | 3 | 1.3 | 3 | 6 | 5 | 2.2 | 5 | 10 | 7 | 3.1 | 2.3 |
| UL21 | 1,599 | 533 | 4 | 3 | 0.6 | 3 | 3 | 2 | 0.4 | 2 | 11 | 7 | 1.3 | 1.8 |
| UL22 | 2,517 | 839 | 22 | 7 | 0.8 | 0.5 | 20 | 6 | 0.7 | 0.4 | 46 | 18 | 2.1 | 0.6 |
| UL23 | 1,131 | 377 | 7 | 5 | 1.3 | 2.5 | 11 | 6 | 1.6 | 1.2 | 20 | 12 | 3.2 | 1.5 |
| UL24 | 846 | 282 | 12 | 3 | 1.1 | 0.3 | 14 | 4 | 1.4 | 0.4 | 26 | 11 | 3.9 | 0.7 |
| UL25 | 1,758 | 586 | 7 | 3 | 0.5 | 0.8 | 8 | 1 | 0.2 | 0.1 | 25 | 4 | 0.7 | 0.2 |
| UL26 | 1,914 | 638 | 11 | 3 | 0.5 | 0.4 | 18 | 7 | 1.1 | 0.6 | 36 | 16 | 2.5 | 0.8 |
| UL26.5 | 990 | 330 | 8 | 2 | 0.6 | 0.3 | 11 | 6 | 1.8 | 1.2 | 25 | 14 | 4.2 | 1.3 |
| UL27 | 2,715 | 905 | 16 | 10 | 1.1 | 1.7 | 17 | 8 | 0.9 | 0.9 | 39 | 17 | 1.9 | 0.8 |
| UL28 | 2,358 | 786 | 14 | 7 | 0.9 | 1 | 11 | 5 | 0.6 | 0.8 | 39 | 15 | 1.9 | 0.6 |
| UL29 | 3,591 | 1,197 | 27 | 8 | 0.7 | 0.4 | 32 | 11 | 0.9 | 0.5 | 83 | 24 | 2 | 0.4 |
| UL30 | 3,723 | 1,241 | 19 | 12 | 1 | 1.7 | 39 | 20 | 1.6 | 1.1 | 68 | 35 | 2.8 | 1.1 |
| UL31 | 918 | 306 | 6 | 1 | 0.3 | 0.2 | 9 | 2 | 0.7 | 0.3 | 18 | 7 | 2.3 | 0.6 |
| UL32 | 1,797 | 599 | 13 | 4 | 0.7 | 0.4 | 13 | 7 | 1.2 | 1.2 | 20 | 8 | 1.3 | 0.7 |
| UL33 | 393 | 131 | 2 | 0 | 0 | 0 | 1 | 0 | 0 | 0 | 5 | 1 | 0.8 | 0.3 |
| UL34 | 831 | 277 | 8 | 4 | 1.4 | 1 | 7 | 2 | 0.7 | 0.4 | 15 | 5 | 1.8 | 0.5 |
| UL35 | 339 | 113 | 0 | 0 | 0 | 0 | 2 | 1 | 0.9 | 0 | 6 | 4 | 3.5 | 2 |
| UL36 | 9,369 | 3,123 | 65 | 31 | 1 | 0.9 | 73 | 33 | 1.1 | 0.8 | 181 | 82 | 2.6 | 0.8 |
| UL37 | 3,345 | 1,115 | 13 | 8 | 0.7 | 1.6 | 27 | 16 | 1.4 | 1.5 | 54 | 25 | 2.2 | 0.9 |
| UL38 | 1,401 | 467 | 10 | 8 | 1.7 | 4 | 22 | 9 | 1.9 | 0.7 | 38 | 15 | 3.2 | 0.7 |
| UL39 | 3,429 | 1,143 | 64 | 22 | 1.9 | 0.5 | 72 | 25 | 2.2 | 0.5 | 118 | 49 | 4.3 | 0.7 |
| UL40 | 1,014 | 338 | 6 | 2 | 0.6 | 0.5 | 4 | 0 | 0 | 0 | 13 | 3 | 0.9 | 0.3 |
| UL41 | 1,479 | 493 | 6 | 1 | 0.2 | 0.2 | 13 | 2 | 0.4 | 0.2 | 26 | 8 | 1.6 | 0.4 |
| UL42 | 1,413 | 471 | 8 | 4 | 0.8 | 1 | 16 | 8 | 1.7 | 1 | 21 | 12 | 2.5 | 1.3 |

| Gene* | ORF length<br>in NT<sup>^</sup> | protein<br>length (#AA) | 10 Neonatal<br># NT diff.<sup>^</sup> | 10 Neonatal<br># AA<br>changes | 10 Neonatal<br>%AA diff. | 10 Neonatal<br>dN/dS ratio | 10 Adult #<br>NT diff. | 10 Adult #<br>AA changes | 10 Adult<br>%AA diff. | 10 Adult<br>dN/dS ratio | 58 Adult #<br>NT diff. | 58 Adult #<br>AA diff. | 58 Adult<br>%AA diff. | 58 Adult<br>dN/dS ratio |
| --- | --- | --- | --- | --- | --- | --- | --- | --- | --- | --- | --- | --- | --- | --- |
| UL43 | 1,245 | 415 | 10 | 6 | 1.4 | 1.5 | 8 | 6 | 1.4 | 3 | 27 | 17 | 4.1 | 1.7 |
| UL44 | 1,443 | 481 | 13 | 6 | 1.2 | 0.9 | 12 | 7 | 1.5 | 1.4 | 23 | 15 | 3.1 | 1.9 |
| UL45 | 519 | 173 | 1 | 1 | 0.6 | 0 | 3 | 1 | 0.6 | 0 | 6 | 3 | 1.7 | 1 |
| UL46 | 2,169 | 723 | 20 | 9 | 1.2 | 0.8 | 18 | 7 | 1 | 0.6 | 40 | 17 | 2.4 | 0.7 |
| UL47 | 2,091 | 697 | 14 | 6 | 0.9 | 0.8 | 13 | 9 | 1.3 | 2.3 | 43 | 20 | 2.9 | 0.9 |
| UL48 | 1,473 | 491 | 6 | 3 | 0.6 | 1 | 7 | 2 | 0.4 | 0.4 | 18 | 8 | 1.6 | 0.8 |
| UL49 | 903 | 301 | 19 | 7 | 2.3 | 0.6 | 17 | 7 | 2.3 | 0.7 | 30 | 12 | 4 | 0.7 |
| UL49A | 264 | 88 | 3 | 1 | 1.1 | 0.5 | 6 | 2 | 2.3 | 0.5 | 8 | 2 | 2.3 | 0.3 |
| UL50 | 1,110 | 370 | 8 | 4 | 1.1 | 1 | 7 | 4 | 1.1 | 1.3 | 22 | 10 | 2.7 | 0.8 |
| UL51 | 735 | 245 | 6 | 1 | 0.4 | 0.2 | 1 | 0 | 0 | 0 | 11 | 6 | 2.4 | 1.2 |
| UL52 | 3,201 | 1,067 | 22 | 12 | 1.1 | 1.2 | 31 | 11 | 1 | 0.6 | 54 | 25 | 2.3 | 0.9 |
| UL53 | 1,017 | 339 | 14 | 8 | 2.4 | 1.3 | 16 | 10 | 2.9 | 1.7 | 26 | 14 | 4.1 | 1.2 |
| UL54 | 1,539 | 513 | 5 | 0 | 0 | 0 | 7 | 3 | 0.6 | 0.8 | 24 | 12 | 2.3 | 1 |
| UL55 | 561 | 187 | 3 | 1 | 0.5 | 0.5 | 2 | 2 | 1.1 | 0 | 9 | 3 | 1.6 | 0.5 |
| UL56 | 708 | 236 | 6 | 4 | 1.7 | 2 | 8 | 5 | 2.1 | 1.7 | 19 | 11 | 4.7 | 1.4 |
| RL2 | 2,502 | 834 | 43 | 22 | 2.6 | 1 | 33 | 19 | 2.3 | 1.4 | 48 | 23 | 2.8 | 0.9 |
| US1 | 1,245 | 415 | 16 | 5 | 1.2 | 0.5 | 14 | 6 | 1.4 | 0.8 | 38 | 22 | 5.3 | 1.4 |
| US2 | 876 | 292 | 7 | 4 | 1.4 | 1.3 | 4 | 3 | 1 | 3 | 15 | 7 | 2.4 | 0.9 |
| US3 | 1,446 | 482 | 9 | 4 | 0.8 | 0.8 | 7 | 4 | 0.8 | 1.3 | 17 | 4 | 0.8 | 0.3 |
| US4 | 2,100 | 700 | 22 | 13 | 1.9 | 1.4 | 40 | 25 | 3.6 | 1.7 | 70 | 45 | 6.4 | 1.8 |
| US5 | 279 | 93 | 4 | 1 | 1.1 | 0.3 | 8 | 4 | 4.3 | 1 | 11 | 6 | 6.5 | 1.2 |
| US6 | 1,182 | 394 | 7 | 2 | 0.5 | 0.4 | 8 | 2 | 0.5 | 0.3 | 18 | 5 | 1.3 | 0.4 |
| US7 | 1,119 | 373 | 10 | 6 | 1.6 | 1.5 | 11 | 5 | 1.3 | 0.8 | 27 | 14 | 3.8 | 1.1 |
| US8 | 1,638 | 546 | 12 | 9 | 1.6 | 3 | 9 | 9 | 1.6 | 0 | 28 | 20 | 3.7 | 2.5 |
| US8A | 441 | 147 | 4 | 3 | 2 | 3 | 2 | 0 | 0 | 0 | 7 | 2 | 1.4 | 0.4 |
| US9 | 270 | 90 | 2 | 0 | 0 | 0 | 3 | 1 | 1.1 | 0.5 | 7 | 4 | 4.4 | 1.3 |
| US10 | 261 | 87 | 7 | 1 | 1.1 | 0.2 | 3 | 2 | 2.3 | 2 | 6 | 2 | 2.3 | 0.5 |
| US11 | 456 | 152 | 2 | 2 | 1.3 | 0 | 0 | 0 | 0 | 0 | 136 | 5 | 3.3 | 0 |
| US12 | 909 | 303 | 6 | 4 | 1.3 | 2 | 6 | 0 | 0 | 0 | 11 | 10 | 3.3 | 10 |

\* See **Table S1** for list of strain names and genome accessions for comparisons of 10 neonatal, 10 adult, and 58 adult HSV-2 genomes.
<sup>^</sup> Abbreviations: ORF, open reading frame; NT, nucleotide; AA, amino acid; diff., difference; %, percent; dN/dS, ratio of nonsynonymous to synonymous nucleotide changes

**Table S3.**
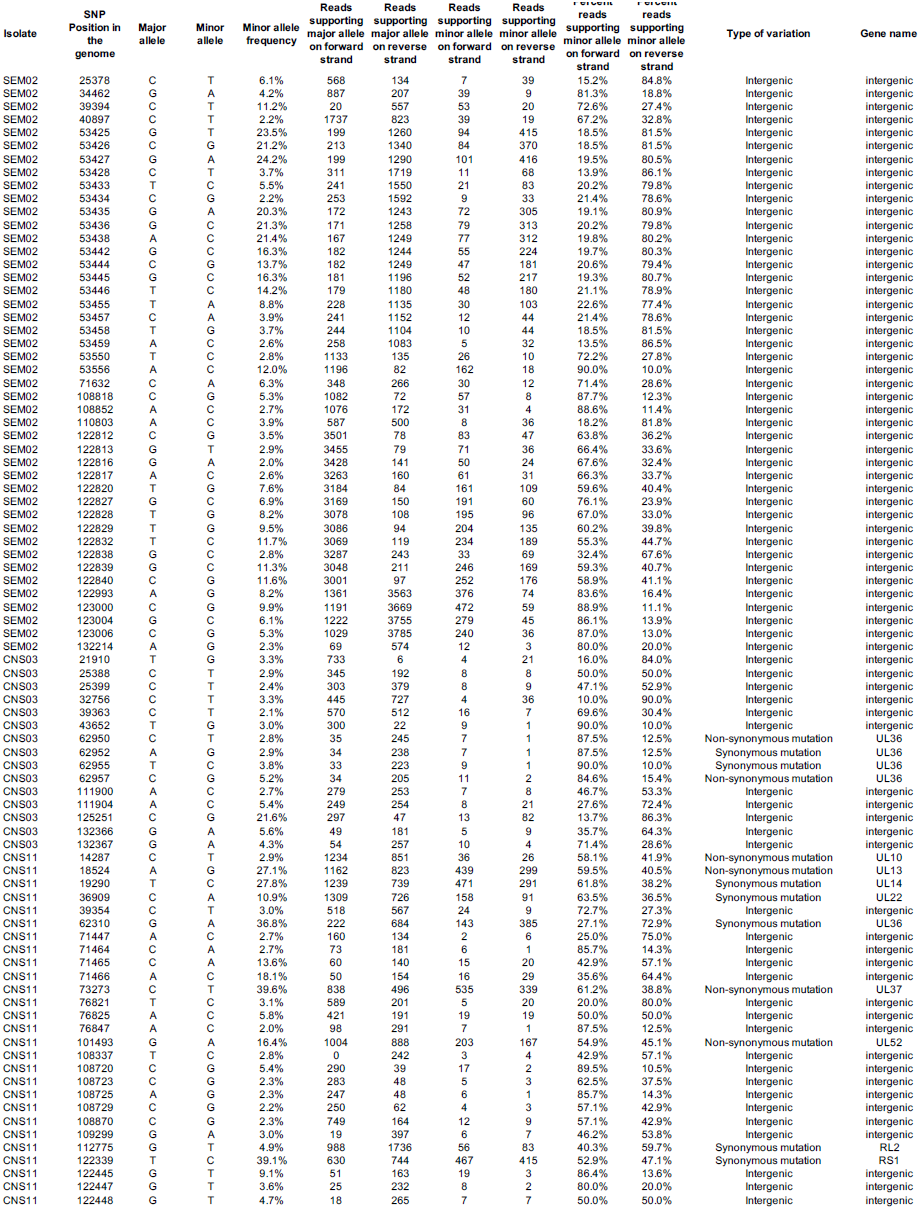
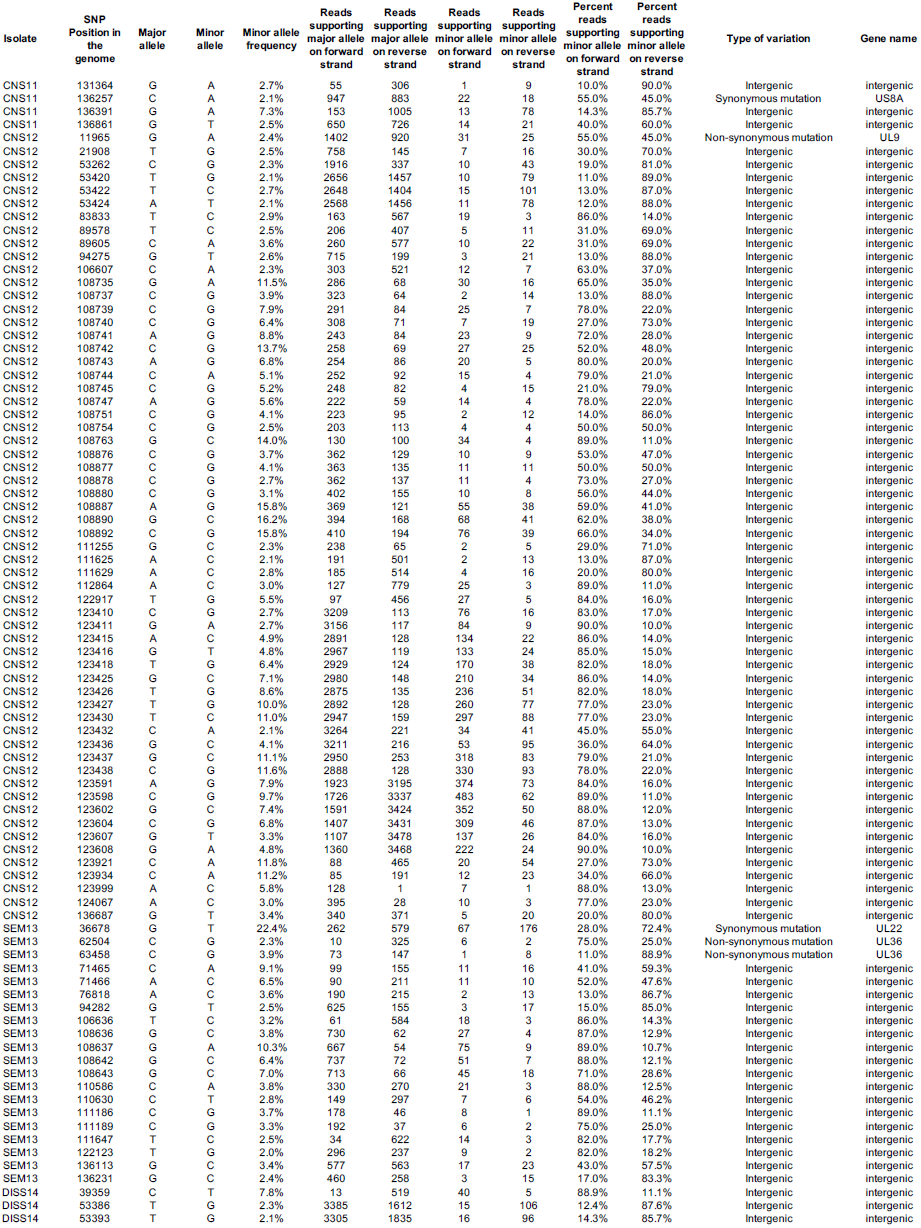
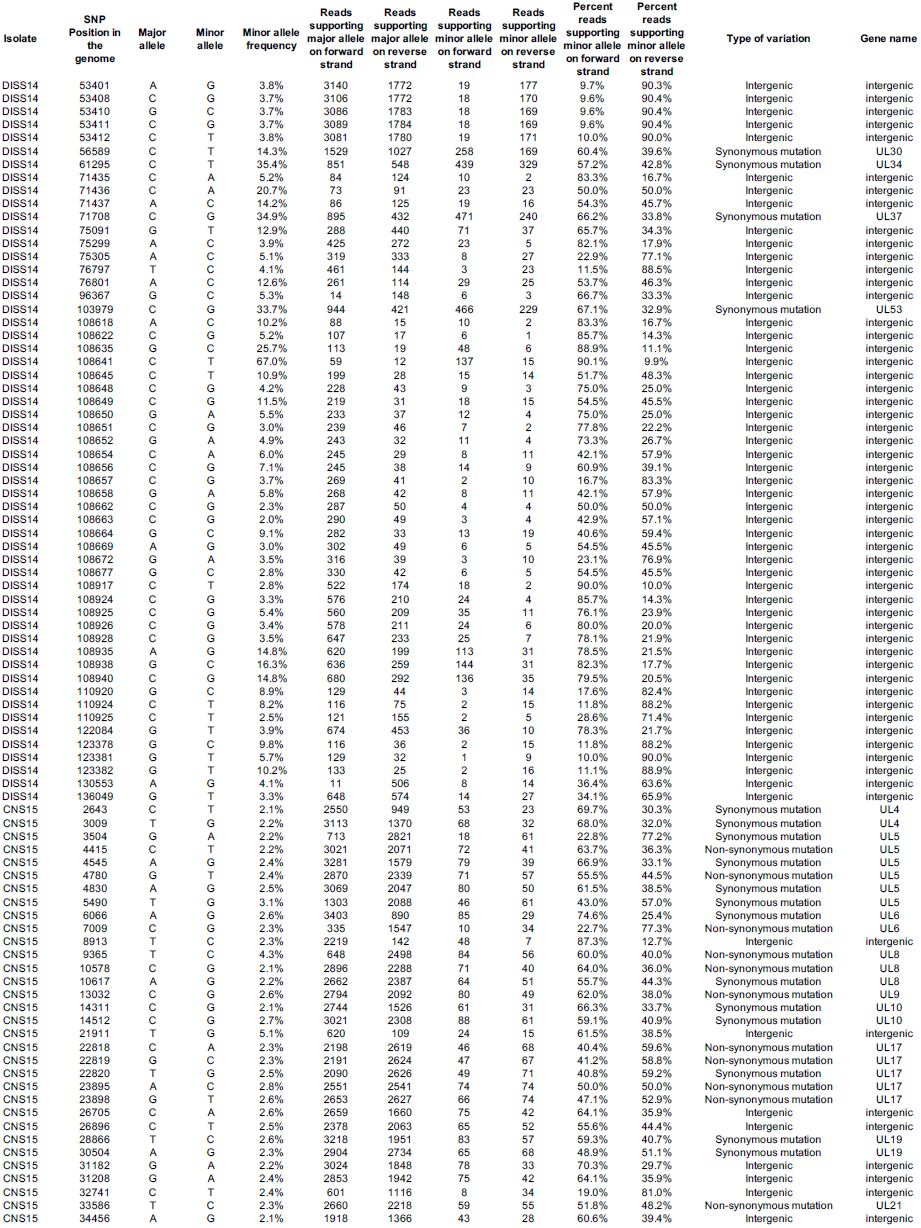
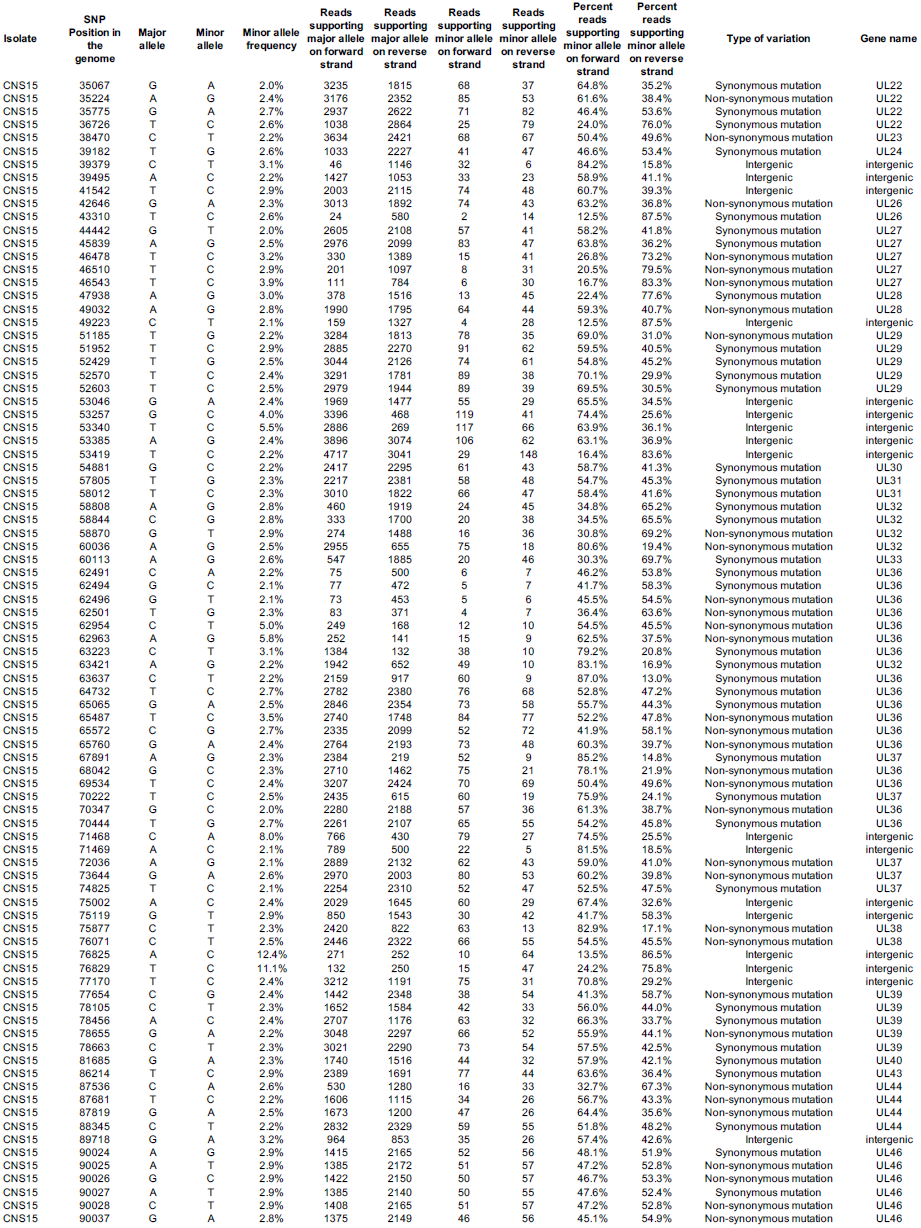
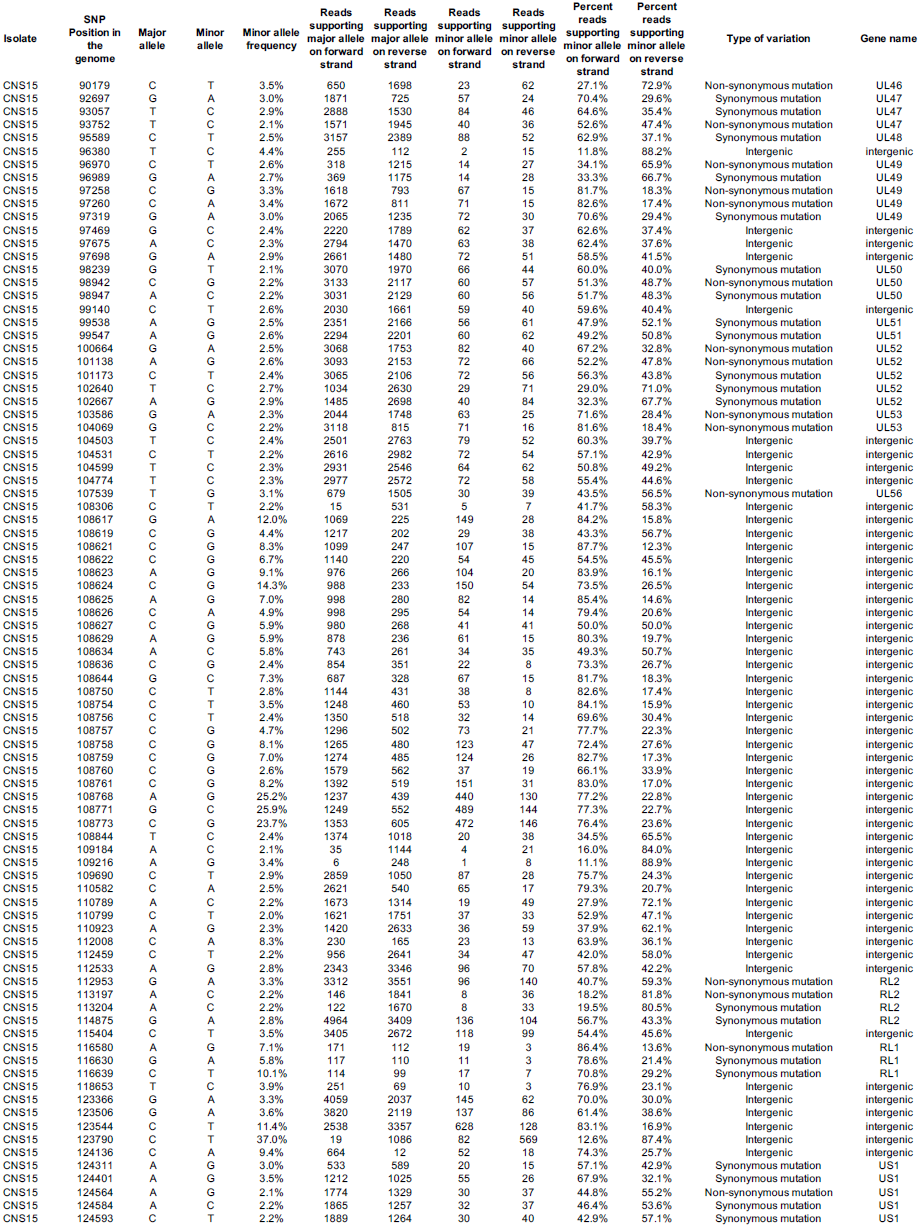
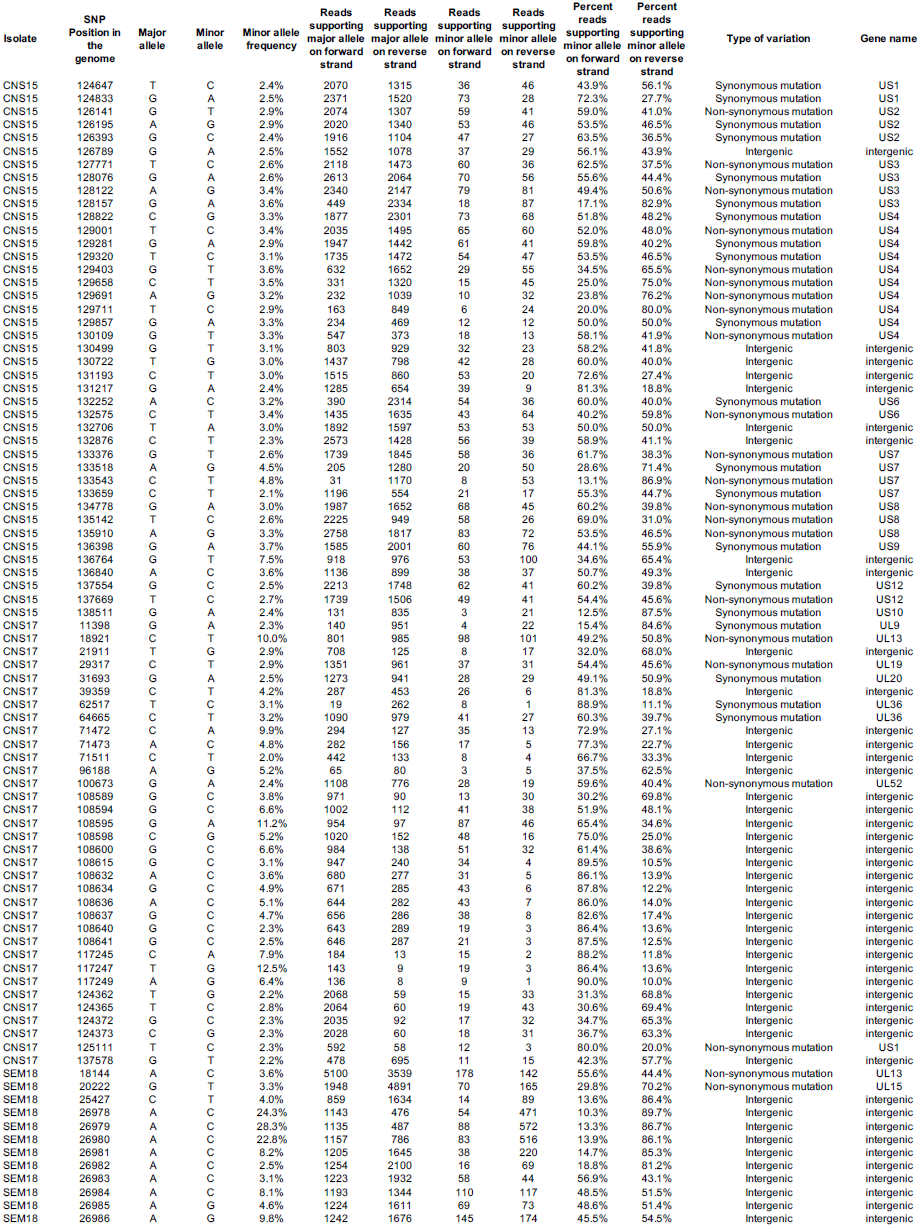
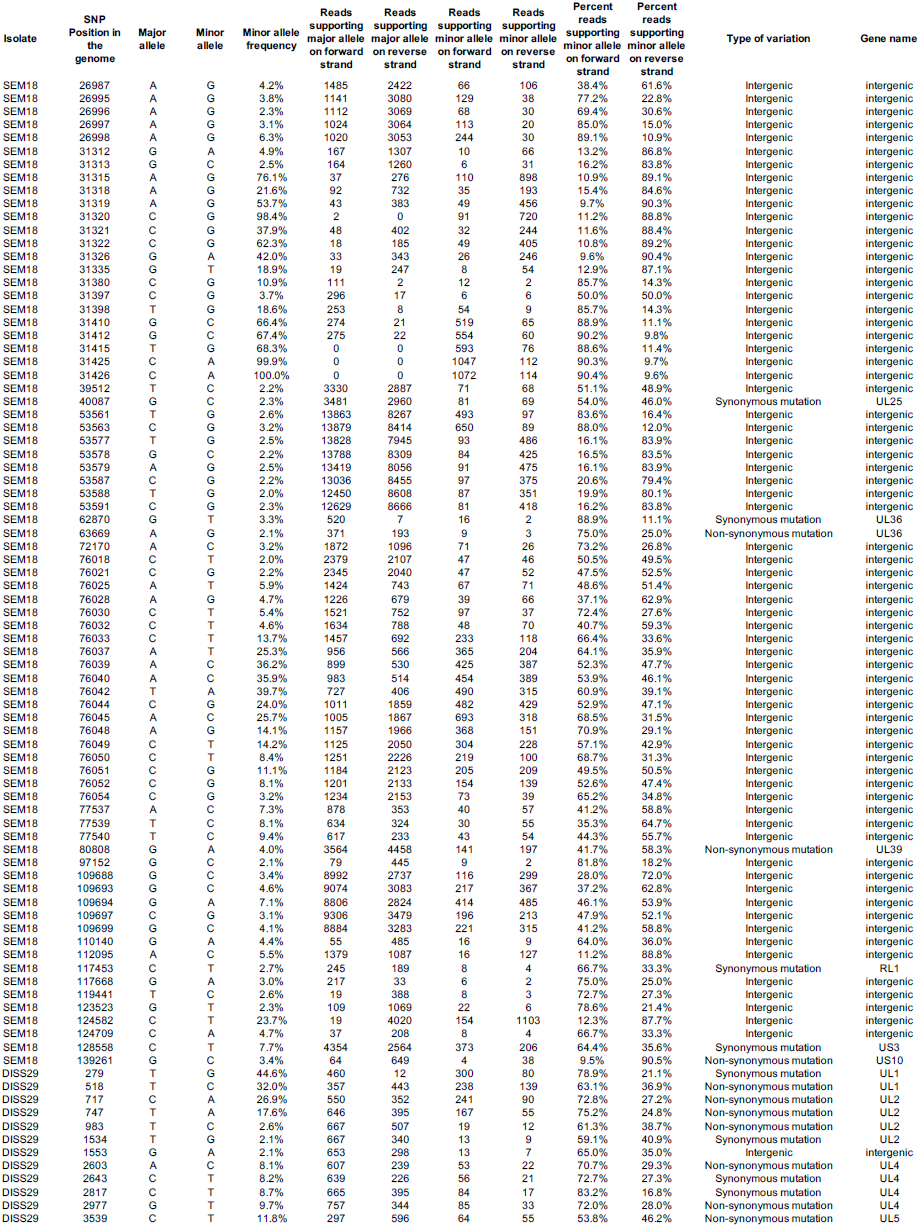
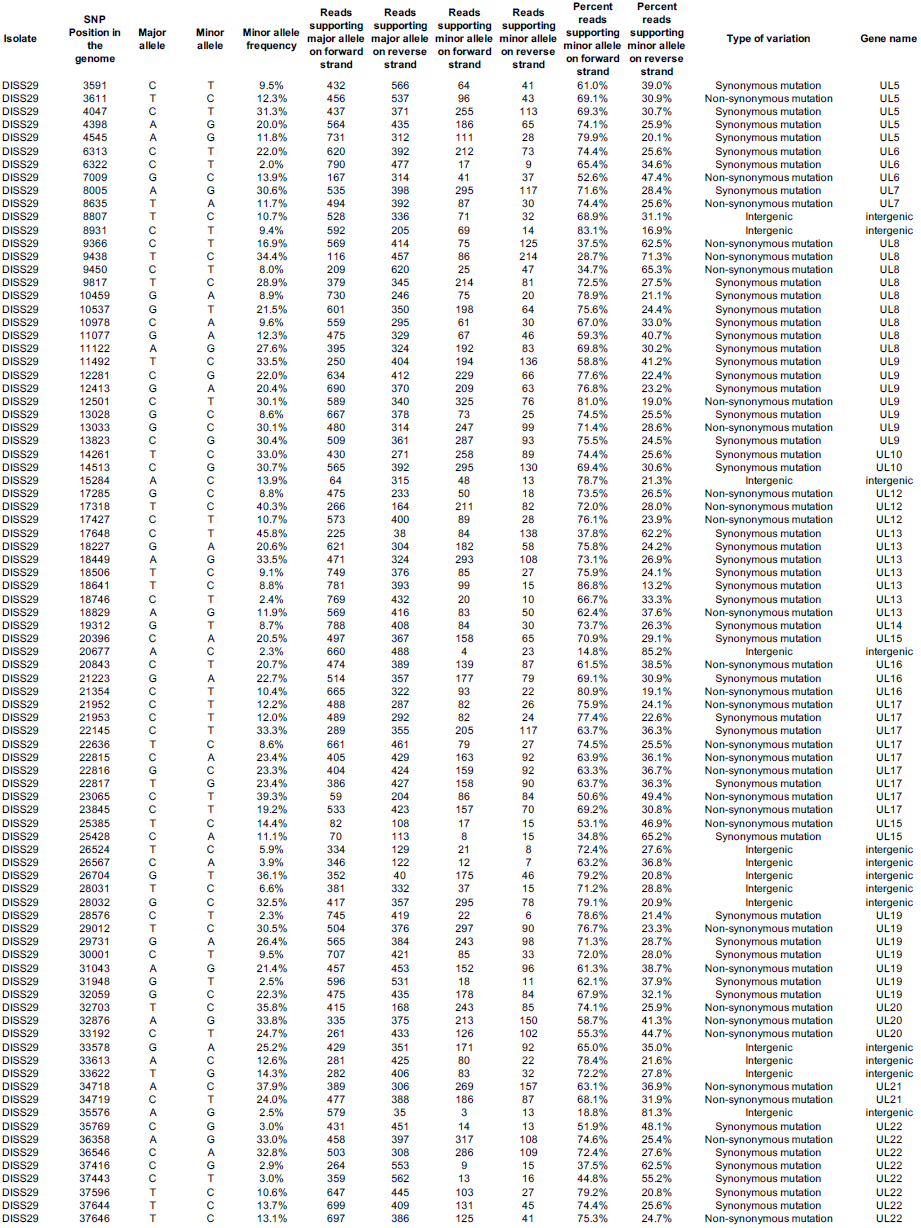
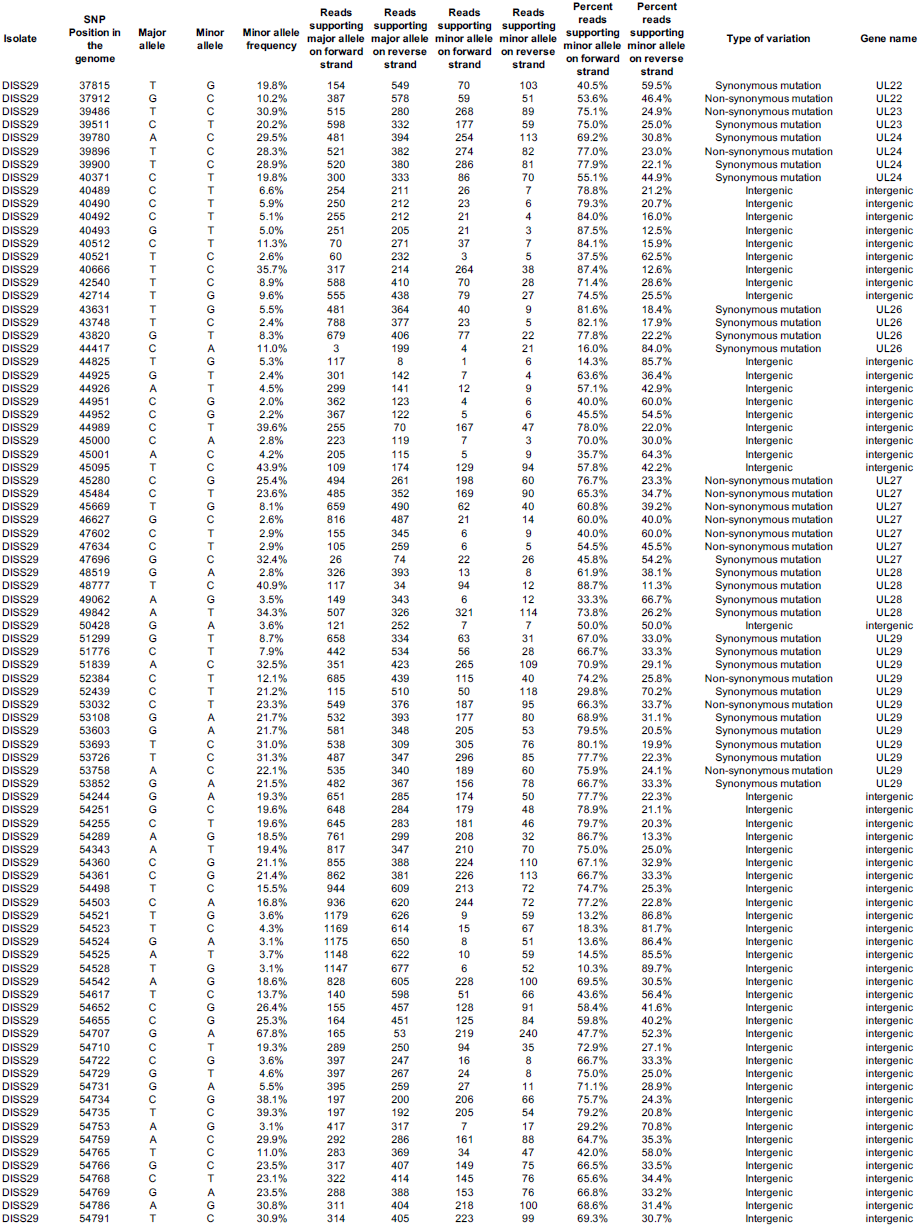
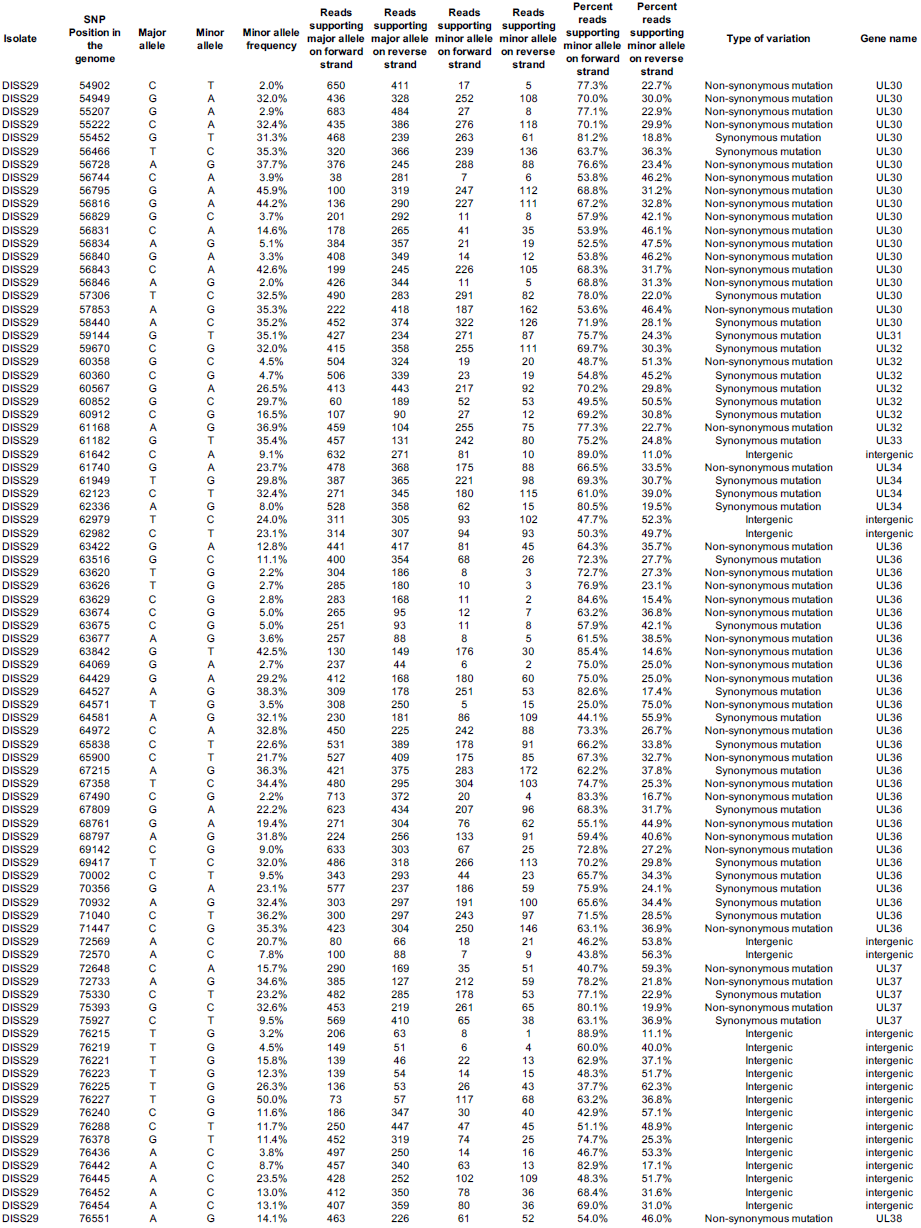
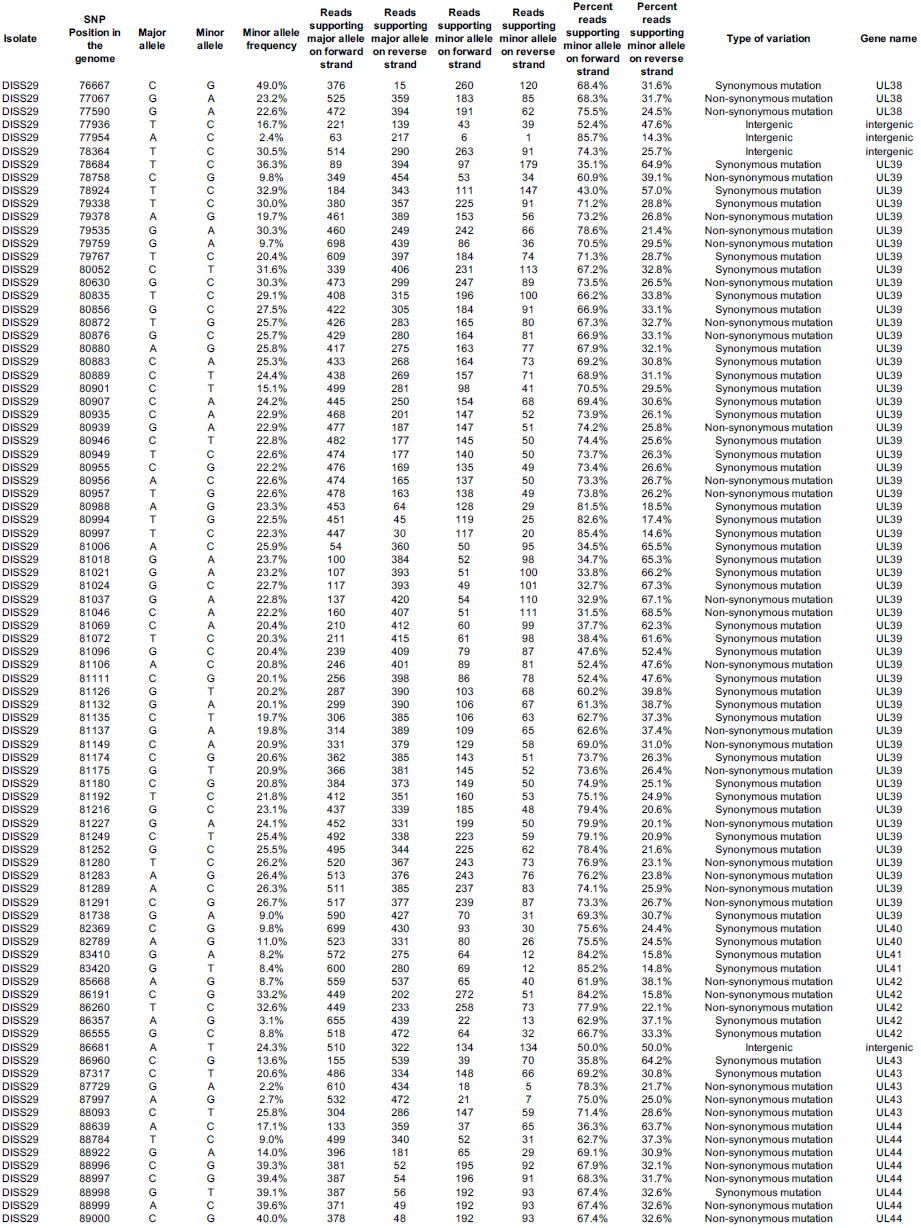
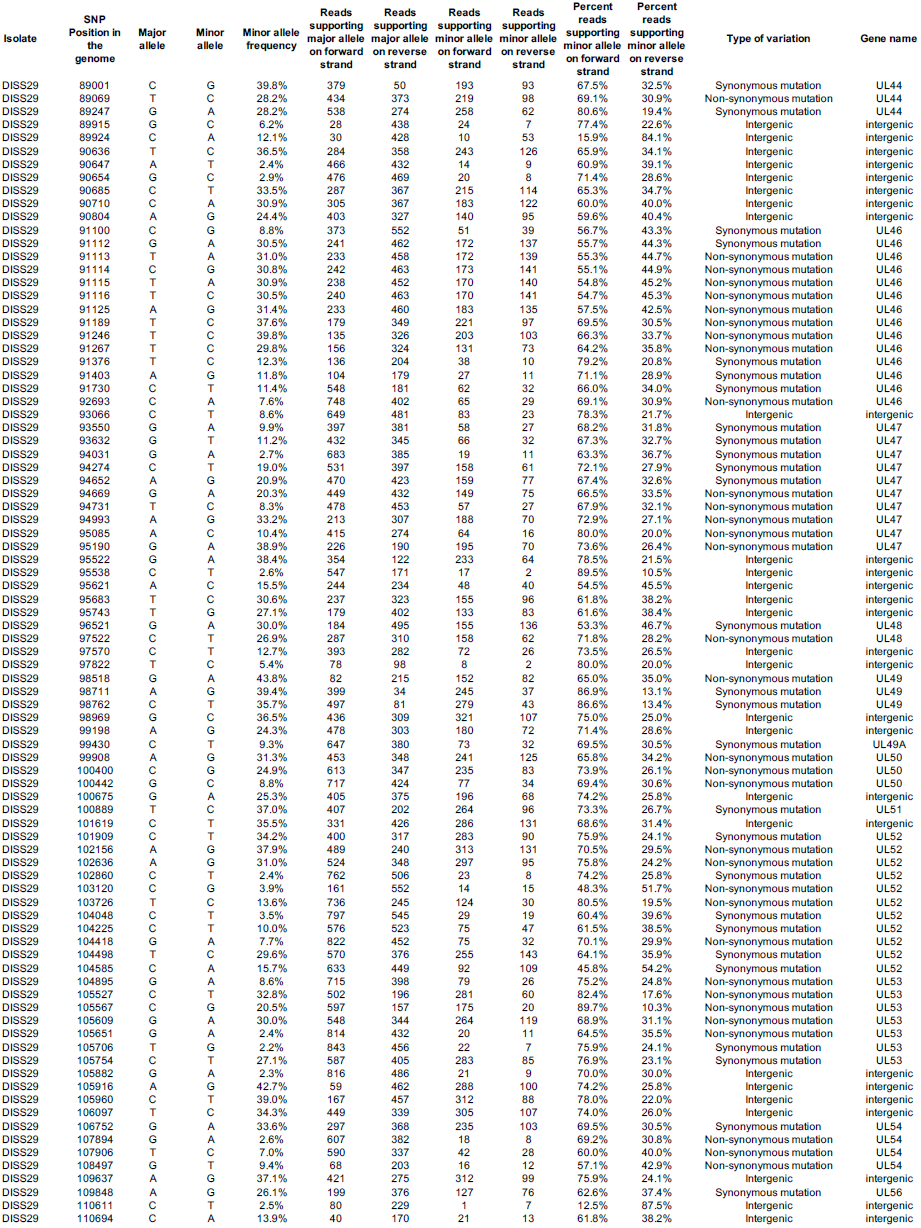
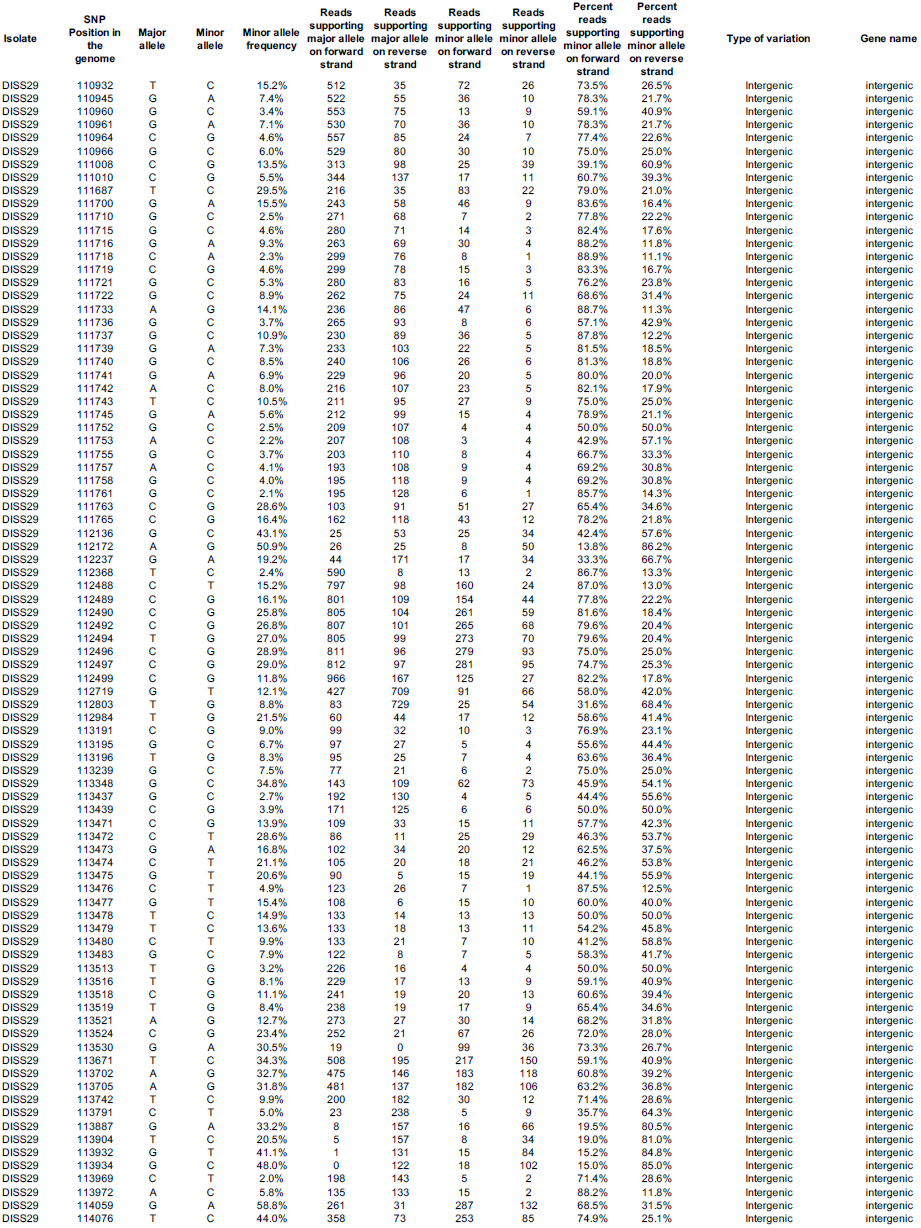
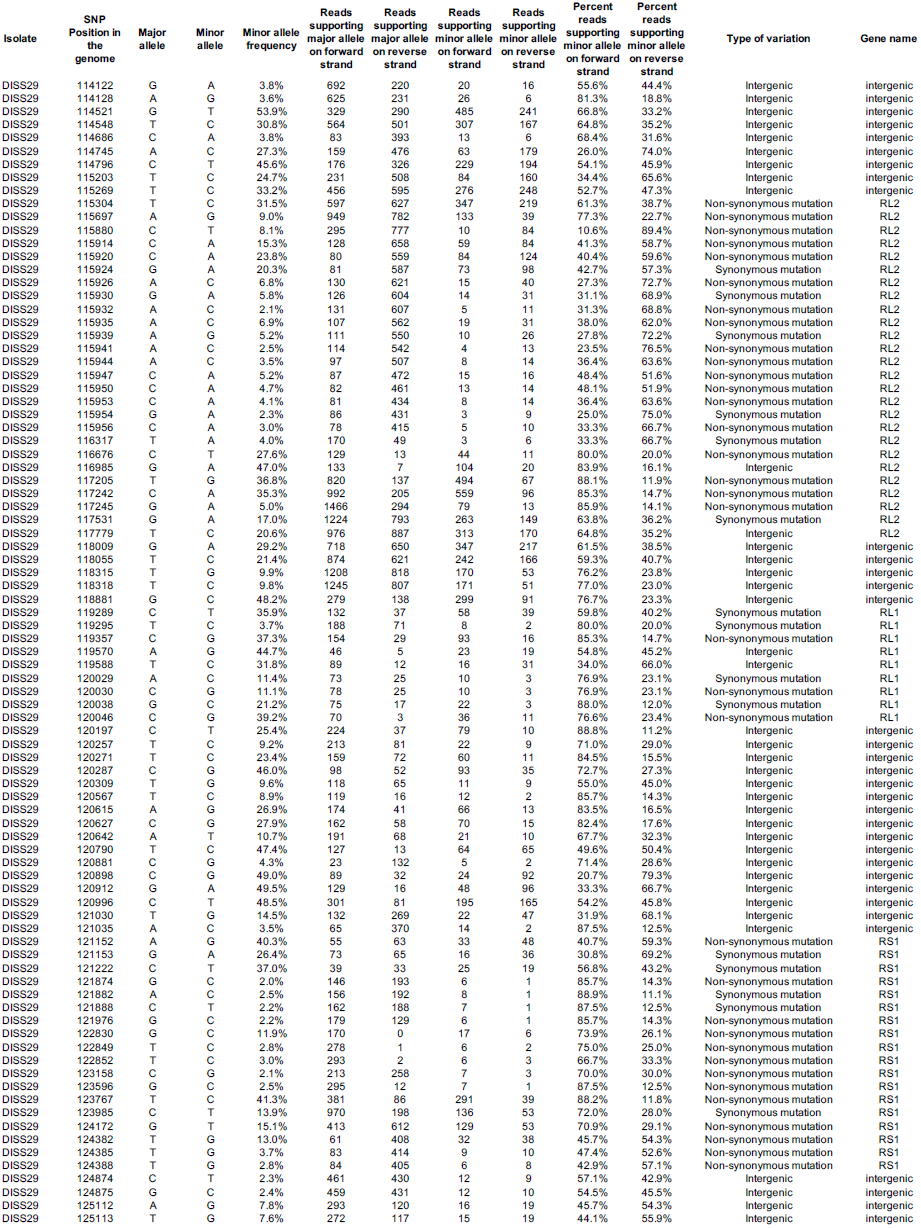
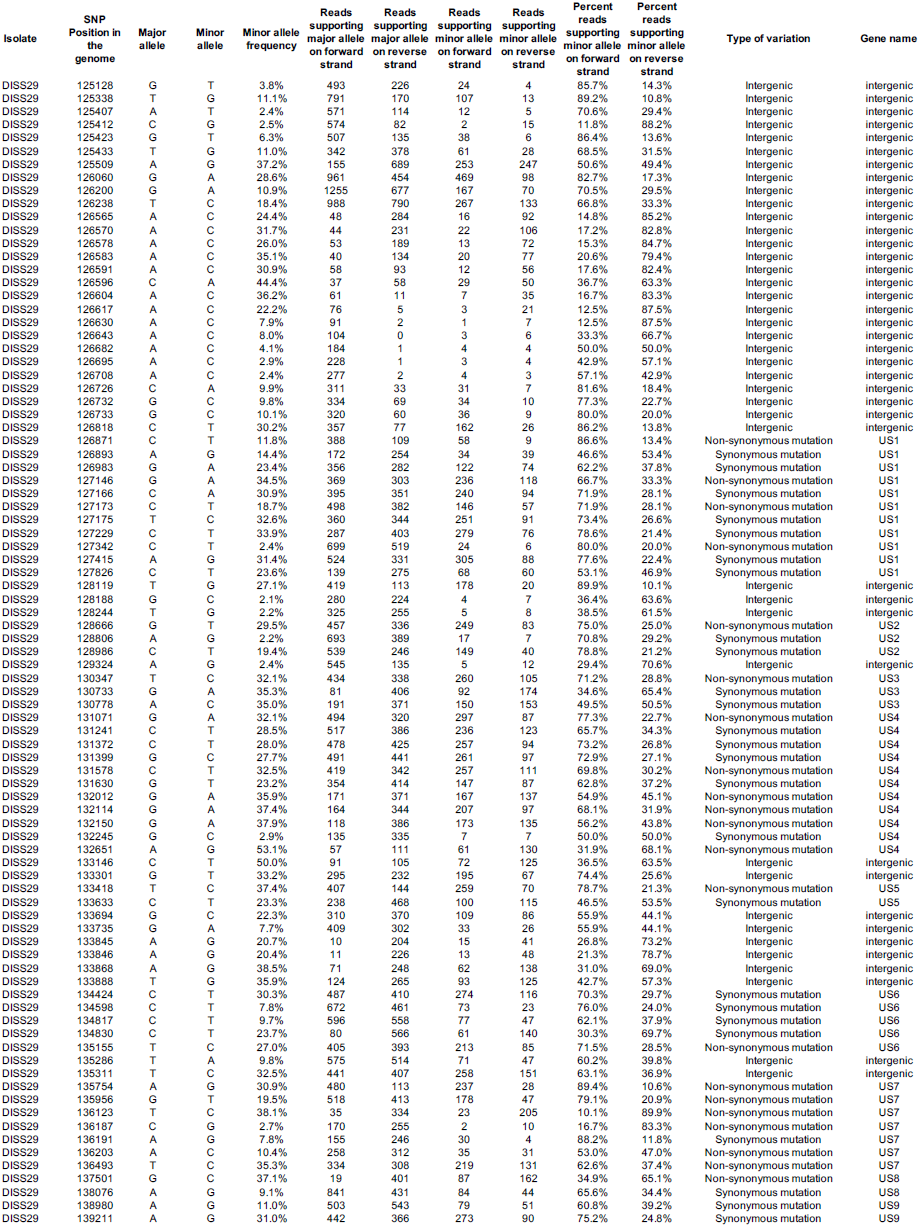
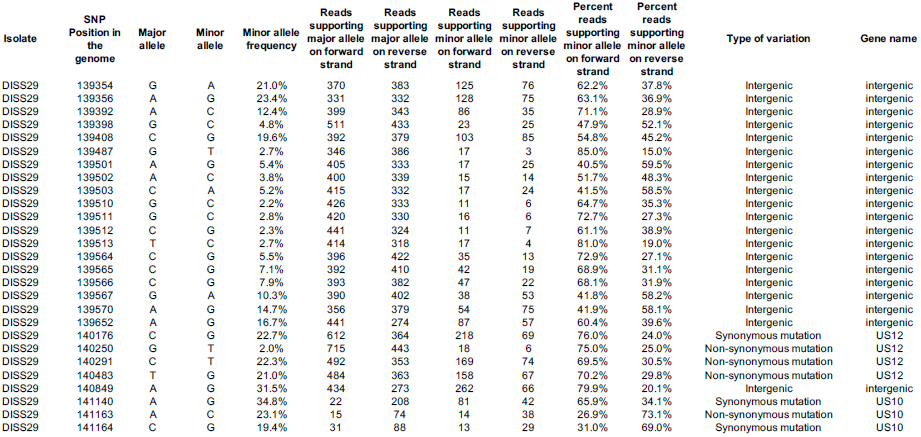
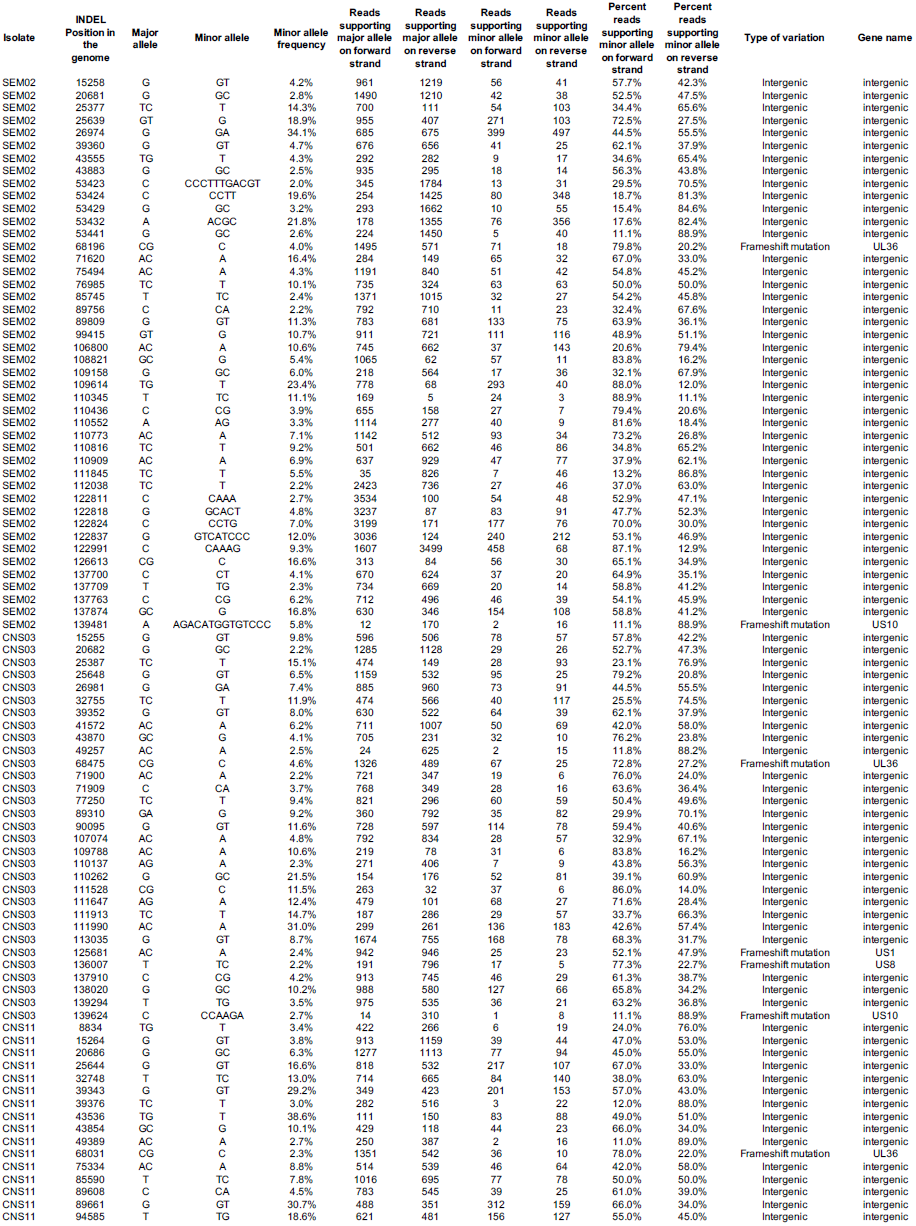
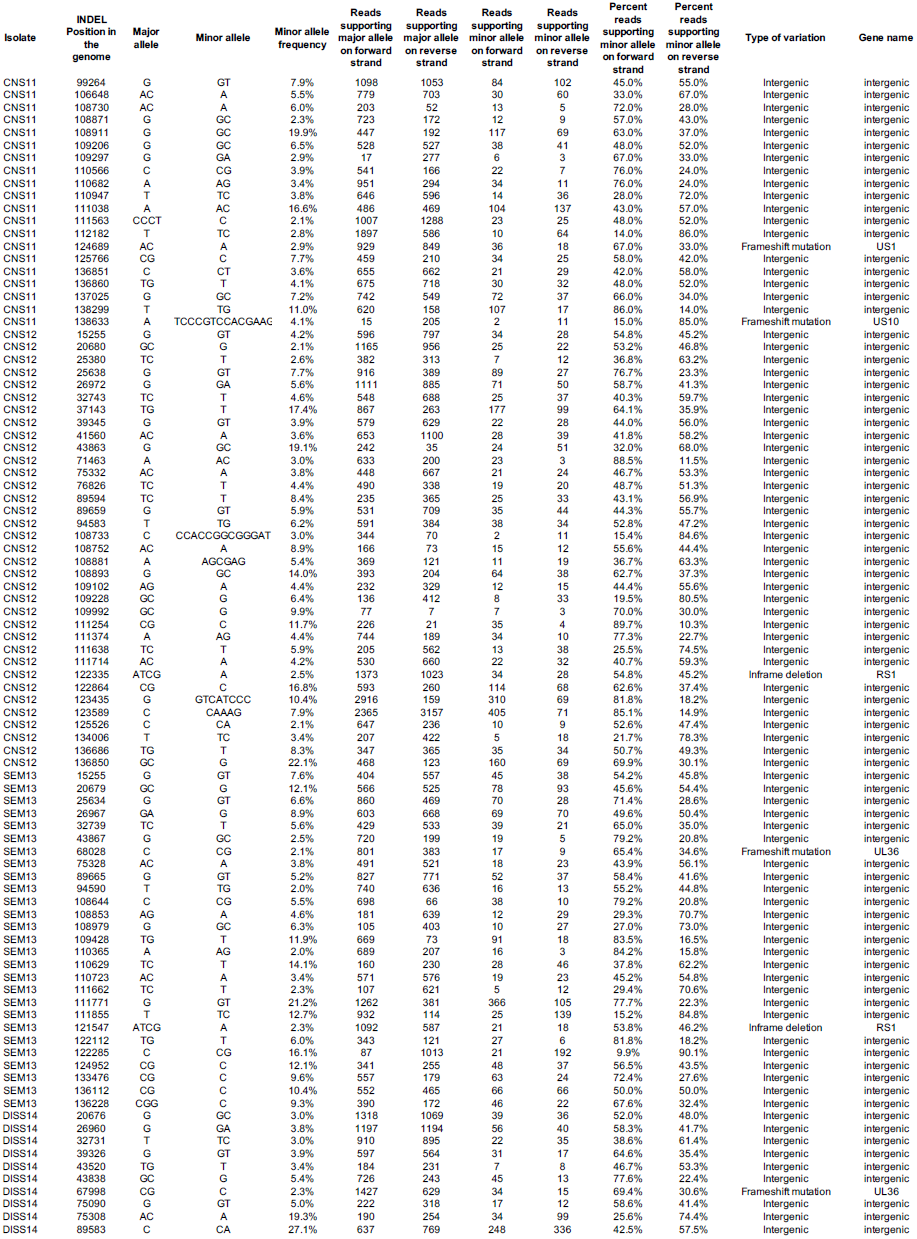
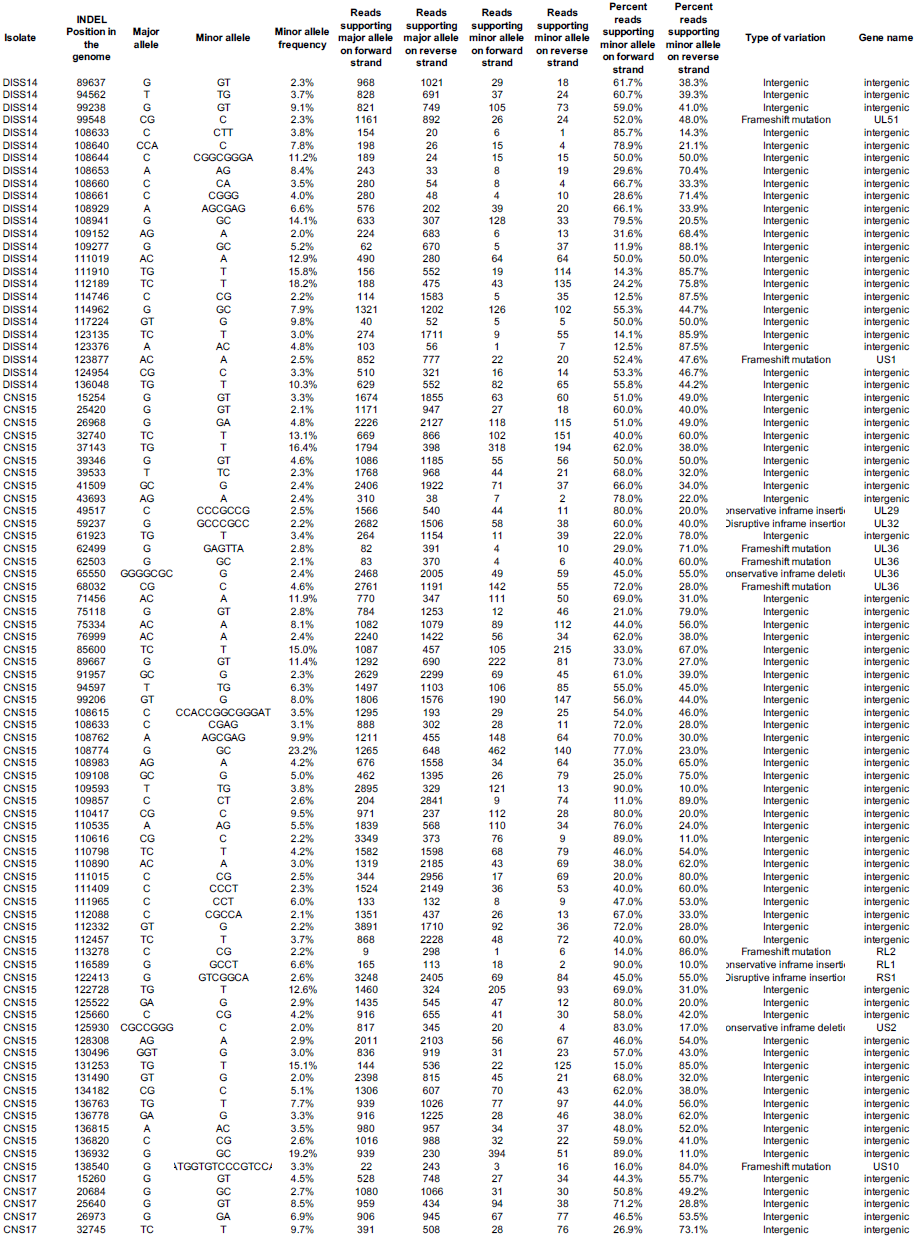
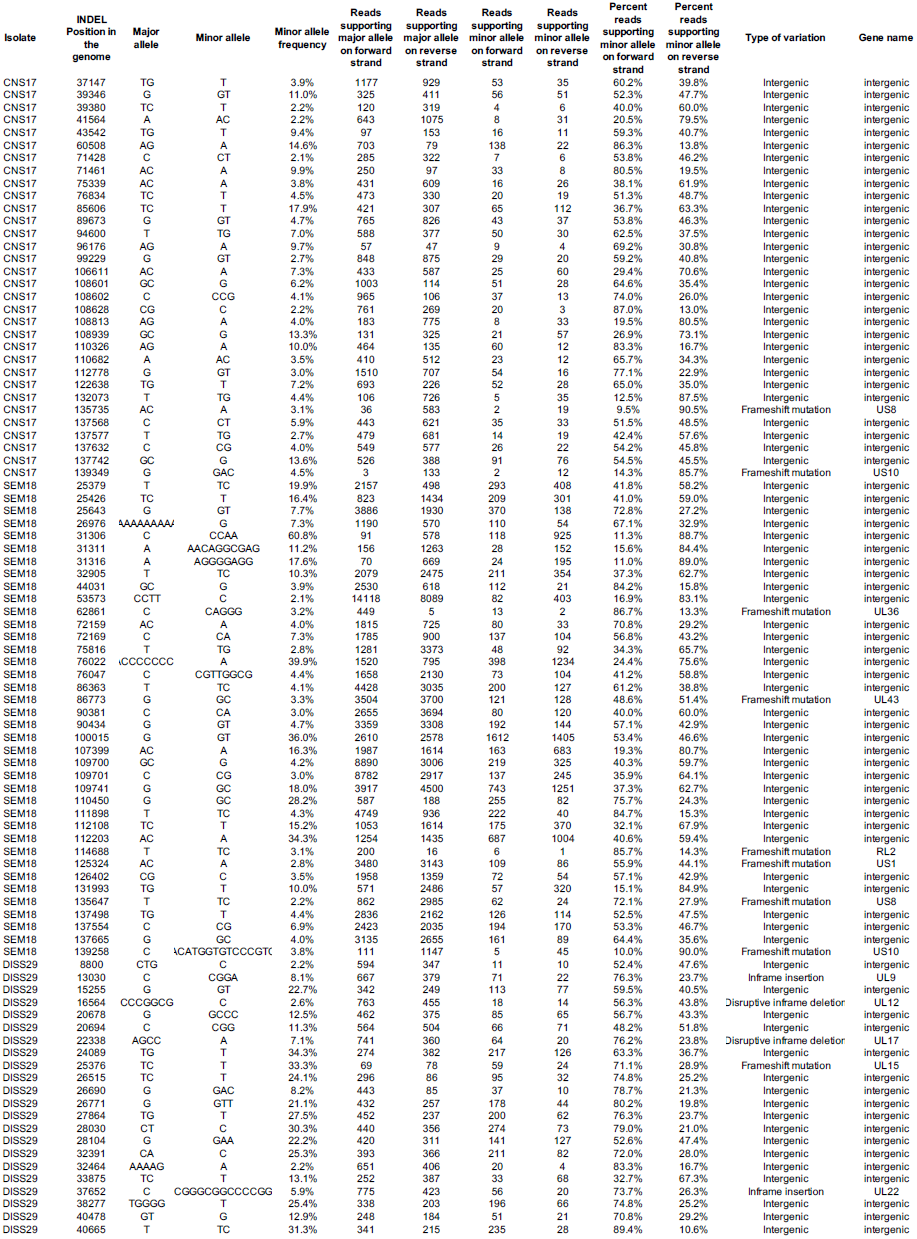
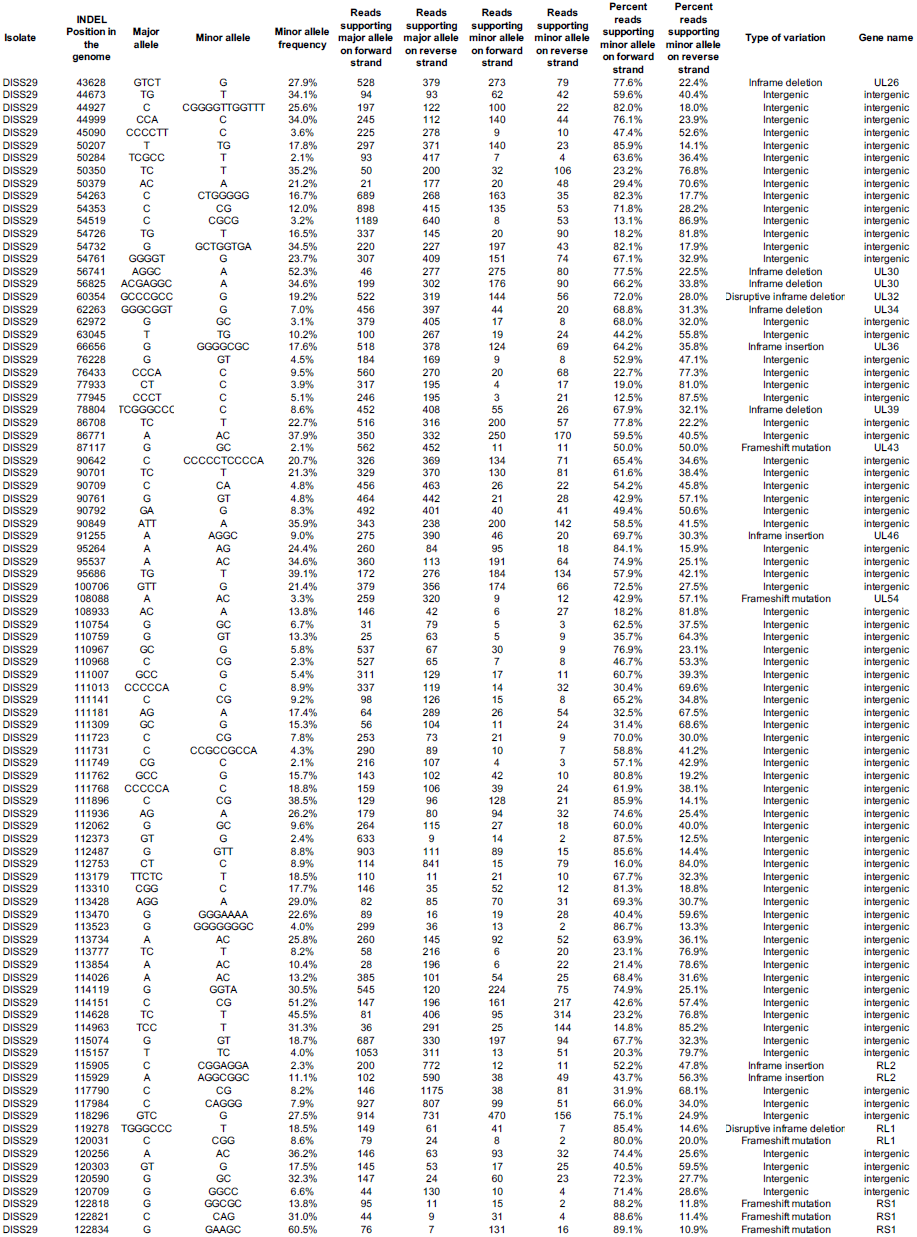
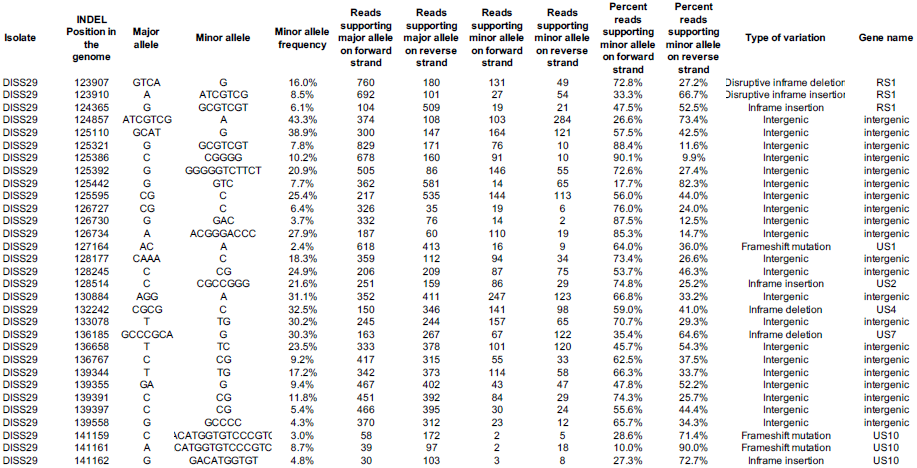
Minor variants -- SNPs and INDELS -- detected in neonatal HSV-2 genomes (two Excel tabs)

**Supplemental Table S4.** Viral genetic variations potentially associated with clinical groups.

| **HSV-2 Gene: Variation** | **Clinical Group (Isolates w/ Variation)** | **Predicted effect<sup>^</sup>** | **Protein Function** | **Neurovirulence** |
| --- | --- | --- | --- | --- |
| **gK (UL53):**<br>V323M | All CSF Isolates (CNS11, DISS14) | Loss of one TM domain | Glycoprotein at virion and cell surface; interacts with gB and UL20 (1); important for cytoplasmic envelopment, viral egress, and virus-induced cell fusion | gK null virus exhibits poor neuronal spread in culture including decreased retrograde and anterograde transport (2); reduced spread within eye and to nervous system, as well as increased survival, following ocular challenge in mice (3) |
| **gI (US7):**<br>R159L + P215S | All CSF Isolates (Both: CNS11,DISS14<br><u>R159L only:</u> CNS03, SEM13,CNS15*,DISS29*<br><u>P215S only:</u> CNS15*,DISS29) | R159L: adds one TM domain;<br>P215S: no change | Glycoprotein at virion and cell surface; forms heterodimer with gE to form viral Fc receptor (4); required for cell-to-cell spread of virus in epithelial and neuronal cells (5, 6); | gI null virus exhibits poor neuronal spread in culture; reduced spread in retina and to retinorecipient areas of the brain following ocular challenge in mice (6) |
| **UL8:** R221S | All CSF Isolates (CNS11, DISS14) | No predicted change | Putative primase subunit of helicase primase complex; required for unwinding of viral DNA | None identified |
| **US2:** F137L | All CSF Isolates (CNS11, DISS14) | No predicted change | Tegument protein; binds and phosphorylates TAK1 to positively regulate NFkB signaling (7) | Virus containing mutated US2 forms small plaques in culture (8); Mutant virus containing a deletion of both US2 and PK result in decreased death following intracerebral injection in mice (9) |
| **gG (US4):**<br>R338L,S442P, E574D | All CSF Isolates (CNS11, DISS14) | No predicted change | Binds and enhances function of chemokines (10); enhances neurotrophin-dependent axonal growth in culture (11) | Virus containing mutated gG results in decreased death following intracerebral injection in mice (8) |
| **UL24:** V93A | No CNS Isolates; only found in SEM isolates (SEM02, SEM13, SEM18) | No predicted change | Membrane-associated nuclear protein; associated with dispersal of nucleolin during infection (12, 13) | UL24 null virus forms small plaques in culture (14); decreased viral load in eye and nervous system following ocular challenge in mice (15–17) |
| **UL20:** P129L | All DISS Isolates w/ CNS Involved (DISS14, DISS29) | Change in position of three TM domains | Membrane-associated protein interacts with gK and gB (1); required for gK glycosylation, cell surface expression (18), and virus-induced cell fusion (19); important for viral egress (20) | UL20 null virus forms small plaques in culture (19); RNA inhibition of UL20 results in decreased rates of encephalitis following HSV footpad infection in mice (21) |

\* present as minor variant
<sup>^</sup> Prediction by SMART Analysis

